# The disordered N-terminal tail of SARS CoV-2 Nucleocapsid protein forms a dynamic complex with RNA

**DOI:** 10.1101/2023.02.10.527914

**Authors:** Jasmine Cubuk, Jhullian J. Alston, J. Jeremías Incicco, Alex S. Holehouse, Kathleen B Hall, Melissa D. Stuchell-Brereton, Andrea Soranno

## Abstract

The SARS-CoV-2 Nucleocapsid (N) protein is responsible for condensation of the viral genome. Characterizing the mechanisms controlling nucleic acid binding is a key step in understanding how condensation is realized. Here, we focus on the role of the RNA Binding Domain (RBD) and its flanking disordered N-Terminal Domain (NTD) tail, using single-molecule Förster Resonance Energy Transfer and coarse-grained simulations. We quantified contact site size and binding affinity for nucleic acids and concomitant conformational changes occurring in the disordered region. We found that the disordered NTD increases the affinity of the RBD for RNA by about 50-fold. Binding of both nonspecific and specific RNA results in a modulation of the tail configurations, which respond in an RNA length-dependent manner. Not only does the disordered NTD increase affinity for RNA, but mutations that occur in the Omicron variant modulate the interactions, indicating a functional role of the disordered tail. Finally, we found that the NTD-RBD preferentially interacts with single-stranded RNA and that the resulting protein:RNA complexes are flexible and dynamic. We speculate that this mechanism of interaction enables the Nucleocapsid protein to search the viral genome for and bind to high-affinity motifs.

## INTRODUCTION

The SARS-CoV-2 virus is a positive-sense single-stranded RNA coronavirus with a genome of nearly 30000 nucleotides^[1]^. This large genome is packaged into small viral particles of ∼100 nm diameter^[2]^. Such a degree of packaging is mediated by the interaction of the viral genome with multiple copies of the Nucleocapsid (N) protein. The “beads on a string structures”^[3,4]^ formed by the SARS-CoV-2 N protein inside the virion are at variance with previously proposed helical structures seen in other coronaviruses^[5,6]^ and the mechanism of their formation is not well understood. From a biophysical standpoint, the compaction of a single viral genome and the phase separation of the protein with multiple nucleic acids potentially stem from the same set of interactions^[7]^. Independent experiments from many labs (including ours) have demonstrated that N protein can undergo phase separation with nucleic acid, both *in vitro* and in living cells^[8–16]^. Phase separation can be favored by specific RNA sequence motifs^[10]^ and altered, in cells, by interactions with small molecules^[17]^. Quantifying the molecular interactions at play is therefore key to identifying the processes controlling condensation on the single- and multi-chain scale.

The SARS-CoV-2 N protein shares a similar domain architecture to analogous N proteins from other coronaviruses, including an RNA Binding Domain (RBD), a dimerization domain, and three intrinsically disordered regions (IDRs) that flank the folded domains. By combining single-molecule experiments and Monte Carlo simulations, we previously showed that N protein adopts a complex and dynamic conformational ensemble as a result of its disordered regions^[7]^. While many experiments have focused on the interaction of the two folded regions (RBD and dimerization domain) with RNA, little is known about the role played by the three disordered regions in aiding the capture and organization of the nucleic acid. The so-called fly-casting model^[18]^ suggests that IDRs have a larger capture radius compared to rigid proteins, resulting in an amplified recruitment of ligands. At the same time, recent experiments have pointed out the peculiarity of disordered regions in encoding for and modulating binding affinity, showing that complexes of oppositely charged biopolymers may achieve high affinity and retain fast dynamic ensembles^[19]^.

Here, we focused our investigation on the RNA Binding Domain (RBD) of the SARS-CoV-2 N protein and studied its interaction with nucleic acids, in the presence and absence of the disordered N-Terminal Domain (NTD). We restricted our analysis to the RBD and the contiguous NTD (**Fig. 1** and **Supplementary Tables 1 and 2**) to identify the specific contributions of the IDR to the folded domain, which otherwise would be masked or altered by the effect of other domains. We hypothesized that the NTD plays an important role since it contributes to localization of the N protein into stress granules^[8,9,20]^ in a RNA dose-dependent manner^[9]^, suggesting that localization is also mediated by its interaction with nucleic acid.

**Figure 1.**
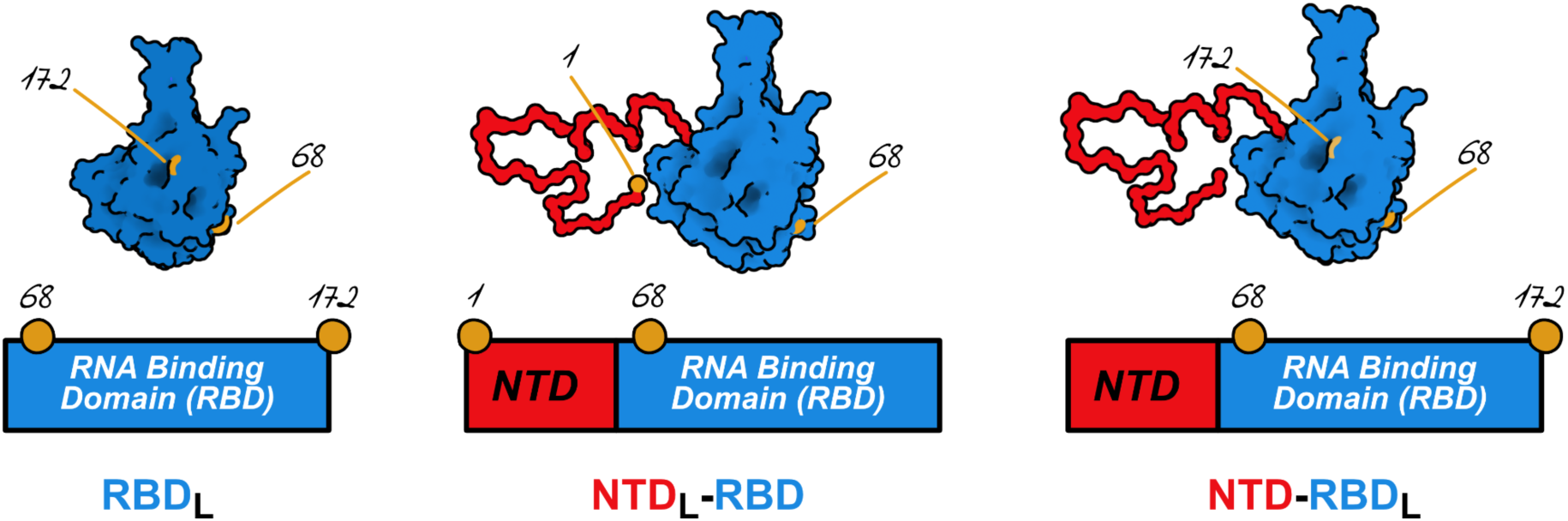
Nucleocapsid protein constructs in this study. (left) RNA Binding Domain (RBD) with dyes in position 68 and 172. (center) NTD-RBD construct with dyes in position 1 and 68, sampling the disordered region. (right) NTD-RBD construct with dyes in position 68 and 172 to sample conformational changes and interactions in the RBD domain.

Single-molecule Förster Resonance Energy Transfer (FRET)^[21–23]^ provides an effective method to determine the affinity and stoichiometry of the binding of RNAs to both RBD and NTD-RBD, while monitoring conformational and dynamic changes occurring in the NTD within the same set of experiments. Single-molecule detection simplifies identification of the contact site size and affinity of the protein even for long nucleic acids since all protein:RNA complexes contain only one single protein (as monitored by Pulsed Interleaved Excitation^[24]^), whereas in typical ensemble experiments one has to account for the contribution of different protein:nucleic acid stoichiometries to the overall signal.

We examined RNA binding using both “nonspecific” and specific RNA molecules. *In cell* crosslinking experiments found that N protein is bound to mRNAs sites containing multiple rU’s^[25]^, while others found it dispersed over the viral genome, comprising both single-stranded and double-stranded regions^[16,26]^. Given the lack of consensus in the literature, we have opted for “nonspecific” poly(rU)_n_ sequences that are well-behaved polyelectrolytes and, differently from poly(rA) and poly(rG), do not undergo stacking at high nucleic acid concentrations. As specific sequences, we have focused on a single-stranded RNA (ssRNA) element of 21 nucleotides that has been isolated from the 5’ UTR of the viral genome (which we will refer to as V21) and on hairpins from the 5’ UTR (SL5B) and a putative packaging signal NSP15^[27]^ (**Supplementary Table 3**).

## MATERIAL AND METHODS

### Protein expression and purification

GST-His9-SARS-CoV2 NTD-RBD_L_ and NTD_L_-RBD Nucleocapsid constructs were expressed recombinantly in Gold BL21(DE3) cells (Agilent). 4 L cultures were grown in LB medium with carbenicillin (100 ug/mL) to OD_600_ ∼0.8 and induced with 0.25 mM IPTG for 3 hours at 37 °C. Harvested cells were lysed with sonication at 4 °C in lysis buffer (50 mM Tris pH 8, 300 mM NaCl, 10% glycerol, 10 mg/mL lysozyme, 5 mM βME, cOmplete™ EDTA-free Protease Inhibitor Cocktail (Roche), DNAse I (NEB), RNAse H (NEB)). The supernatant was cleared by centrifugation (140,000 x g for 1 hr) and bound to a HisTrap FF column (GE Healthcare) in buffer A (50 mM Tris pH 8, 300 mM NaCl, 10% glycerol, 20 mM imidazole, 5 mM βME). The column was then washed with High Salt Buffer (50 mM Tris pH 8, 2M NaCl, 10% glycerol, 5 mM βME) for ten column volumes followed by ten column volumes of Buffer A. GST-His9 -N protein fusion was eluted with buffer B (buffer A + 500 mM imidazole) and dialyzed into cleavage buffer (50 mM Tris pH 8, 50 mM NaCl, 10% glycerol, 1 mM DTT) with HRV 3C protease, thus cleaving the GST-His9 -N fusion yielding N protein with two additional N-term residues (GlyPro). N protein was then bound to an SP sepharose FF column (GE Healthcare) and eluted using a gradient of 0-100% buffer B (buffer A: 50 mM Tris pH 8, 50 mM NaCl, 10% glycerol, 5 mM βME, buffer B: buffer A + 1 M NaCl) over 100 min. Purified NTD-RBD_L_ and NTD_L_-RBD constructs were analyzed using SDS-PAGE and their concentrations were determined spectroscopically in 50 mM Tris (pH 8.0), 500 mM NaCl, 10% (v/v) glycerol using an extinction coefficient of 25200 M^−1^ cm^−1^ at 280 nm.

GST-His9-SARS-CoV2 RBD_L_ Nucleocapsid construct was expressed recombinantly in Gold BL21(DE3) cells (Agilent). 4 L cultures were grown in LB medium with carbenicillin (100 ug/mL) to OD600 ∼ 0.6 and induced with 0.3 mM IPTG for 3 hours at 37 °C. Harvested cells were lysed with sonication at 4 °C in lysis buffer (50 mM Tris pH 7, 300 mM NaCl, 10% glycerol, 10 mg/mL lysozyme, 5 mM βME, cOmplete™ EDTA-free Protease Inhibitor Cocktail (Roche), DNAse I (NEB), RNAse H (NEB)). The supernatant was cleared by centrifugation (140,000 x g for 1 hr) and bound to a HisTrap FF column (GE Healthcare) in buffer A (50 mM Tris pH 7, 300 mM NaCl, 10% glycerol, 20 mM imidazole, 5 mM βME).The column was then washed with High Salt Buffer (50 mM Tris pH 7, 2M NaCl, 10% glycerol, 5 mM βME) for ten column volumes followed by ten column volumes of Buffer A. GST-His9 -N protein fusion was eluted with buffer B (buffer A + 500 mM imidazole) and dialyzed into cleavage buffer (20 mM Tris pH 7, 20 mM NaCl, 10% glycerol, 1 mM DTT) with HRV 3C protease, thus cleaving the GST-His9 -N fusion yielding N protein with two additional N-term residues (GlyPro). The N protein was then run over a HisTrap FF column (GE Healthcare) in Buffer A (20 mM Tris pH 7, 20 mM NaCl, 10% glycerol) and the flow through was collected. N protein was then bound to an SP sepharose FF column (GE Healthcare) and eluted using a gradient of 0-100% buffer B (buffer A: 20 mM Tris pH 7, 50 mM NaCl, 10% glycerol, 5 mM βME, buffer B: buffer A + 1 M NaCl) over 100 min. Purified RBD_L_ construct was analyzed using SDS-PAGE and its concentration was determined spectroscopically in 50 mM Tris (pH 7.0), 300 mM NaCl, 10% (v/v) glycerol using an extinction coefficient of 25200 M^−1^ cm^−1^ at 280 nm. Plasmid DNA sequences for the constructs can be found in **Supplementary Information**.

### Protein labeling

All Nucleocapsid variants were labeled with Alexa Fluor 488 maleimide (Molecular Probes, USA) under denaturing conditions in buffer A (10 mM Tris pH 7.3, 6 M Urea) at a dye/protein molar ratio of 0.7/1 for 2 hrs at room temperature. Single labeled protein was isolated via ion-exchange chromatography (Mono S 5/50 GL, GE Healthcare - protein bound in buffer A (+5 mM βME) and eluted with 0-40% buffer B (buffer A + 1 M NaCl) gradient over 70 min) and UV-Vis spectroscopic analysis to identify fractions with 1:1 dye:protein labeling. Single donor labeled N protein was then subsequently labeled with Alexa Fluor 594 maleimide at a dye/protein molar ratio of 1.3/1 for 2 hrs at room temperature. Double-labeled (488:594) protein was then further purified via ion-exchange chromatography (Mono S 5/50 GL, GE Healthcare).

### RNA preparation

Single-stranded RNAs were purchased from IDT (USA) and Horizon Discovery (USA). Hairpin RNAs were transcribed with T7 RNA polymerase from DNA oligonucleotides (IDT), using T7 RNA polymerase (NEB USA) in an optimized reaction mix. RNAs were purified by denaturing polyacrylamide gel electrophoresis (15% acrylamide, 19:1 bis, 8 M urea, Tris-Borate-EDTA), bands were visualized by UV shadowing and cut out. Gel slices were soaked in 0.3 M sodium acetate overnight at 30 °C in a rotating mixer, the solution was recovered and gel debris removed by centrifugation. RNA was precipitated overnight at −20 °C in the presence of glycogen with 3X volume 100% ethanol, and the pellet resuspended in Milli-Q water (Millipore-Sigma, USA). Hairpins were annealed in 10 mM HEPES pH 6.5, 50 mM KCl buffer and their integrity and stability measured in UV melting experiments as a function of their concentration. RNA concentrations were determined spectrophotometrically employing their computed extinction coefficients at 260 nm.

### Instrumentation

Single-molecule experiments were performed on a modified Picoquant MT200 instrument (Picoquant, Germany) using Pulsed Interleaved Excitation to enable identification of the donor- and acceptor-only as well as donor-acceptor populations. All data reported in this work are selected for the donor-acceptor population. Single-molecule measurements, unless otherwise stated, have been performed in 50 mM Tris, pH 7.4 at room temperature (23 ± 1 °C).

### Analysis of binding experiments

Binding of RNA ligands to labeled N protein constructs was monitored by following either the mean value of the transfer efficiency distribution or the fraction of bursts associated with the bound and unbound population (when they can be resolved).

In the first case, titration curves were analyzed according to:

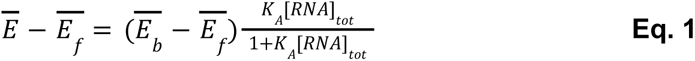

where 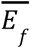 and 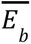 are the mean transfer efficiencies for the free and bound protein, *K*_A_ is the association constant and [RNA] is the total concentration of RNA. Note that under all conditions the free RNA concentration is always much higher than the concentration of a bound complex because of the single-molecule concentrations used in the experiment.

In the second case, when the fraction of bound protein *f_b_* is directly estimated, titration curves were analyzed according to:

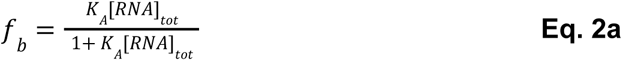

for the 1:1 binding cases, and:

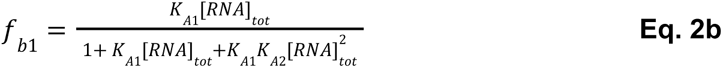

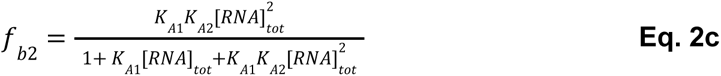

for the 1:2 case treated in this work.

For the special case of the binding to the polynucleotide poly(rU), titration curves were obtained and analyzed as a function of the total concentration of nucleotide residues [poly(rU)], not RNA molecules. This is justifiable because under the experimental conditions employed, where the protein concentration is so much lower than RNA concentration, the McGhee-von Hippel formulation for the binding of large ligands to one dimensional lattices reduces to:

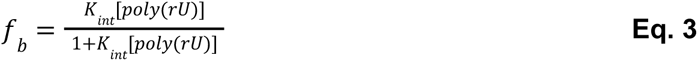

where *K*_int_ is the intrinsic association constant.

### Statistical Analysis

Values associated with multiple measurements are presented as mean and standard deviation of the measured points of at least two points. Results of model fit to the data are presented as best value and corresponding error of the fit as determined using non-linear regression algorithms in Mathematica (Wolfram Research Inc, USA).

### Data Availability, Software, Algorithms

Data analysis of single-molecule data has been performed using the Fretica package for Mathematica (Wolfram Research Inc, USA) developed by the Schuler group (https://schuler.bioc.uzh.ch/wp-content/uploads/2022/07/Fretica20220630.zip). All single-molecule data reported in this work are deposited at https://github.com/holehouse-lab/supportingdata/tree/master/2023/cubuk_2023. Raw photon traces of single-molecule data will be made available upon request.

### Simulations

Coarse-grained molecular dynamics (MD) simulations were performed in the NVT ensemble using the LAMMPS simulation engine with Mpipi model using the default parameters developed by Joseph et al.^[28]^. Mpipi is a one-bead-per residue coarse grained force field developed specifically for working with intrinsically disordered proteins. Non-bonded interactions are driven by a short-range potential and, where applicable, a long-range Coulombic potential. Bonded interactions are encoded via a simple harmonic potential. Simulations were performed with NTD-RBD, RBD, and NTD, with and without (rU)_n_ of lengths n = 10, 12, 15, 17, 20, 25, 30, 35, 40 and 180 nucleotides. We also performed simulations of the Omicron variant of the NTD-RBD, with substitution of the proline residue in position 13 with a leucine (P13L) and deletion of residues from 31 to 33 (Δ^31-33^). For assessing the role of each site, also we performed simulations of the P13L mutation alone versus the Δ^31-33^ alone. All simulations of the Omicron constructs were performed with and without(rU)_25_.

All simulations were run with multiple independent repeats using a 30 nm^3^ simulation box and periodic boundary conditions. As in previous work, folded domains were modeled as rigid bodies, whereas intrinsically disordered regions and ssRNA were described as flexible polymers^[28,29]^. For simulations where folded domains were present (i.e. those with the RBD), six distinct RBD conformations were taken from all-atom simulations of the RBD performed using the Folding@Home distributed computing platform^[7,30,31]^. This enables us to ensure conclusions obtained are not dependent on a specific RBD conformation. For the six independent starting configurations, five repeats were performed, with 300 million steps per repeat, such that 30 independent simulations were run for each unique protein/RNA combination. Simulation configuration data was recorded every 100,000 steps, and the first 600,000 steps (0.2% of the simulation) discarded as equilibration. Across the 30 independent simulations for each protein/RNA combination we generated approximately 270,000 frames. A summary of the simulations performed is provided in **Supplementary Table 4**.

Simulations were analyzed using SOURSOP (https://soursop.readthedocs.io/) and MDTraj ^[32]^. All analysis code for simulations is provided at https://github.com/holehouse-lab/supportingdata/tree/master/2023/cubuk_2023. For more details on the simulations see extended materials and methods in the **Supplementary Information**.

### Database Referencing

Sequence data for the Nucleocapsid variants, including the Omicron variant, were obtained from the GISAID lineage-comparison database: https://gisaid.org/lineage-comparison/

Extended description of experimental procedures, material and methods, and data analysis are presented in **Supplementary Information**.

## RESULTS

In order to investigate the binding and conformational changes of the N-terminal disordered tail and RNA-binding domain of SARS-CoV-2 Nucleocapsid protein *via* single-molecule FRET, we created two truncated constructs, one spanning the full N-terminal segment of the protein comprising both the NTD and RBD and another comprising the RBD alone (**Fig. 1**). Cysteine mutations were introduced in the wild-type sequence to enable fluorophore addition to the constructs *via* maleimide-thiol chemistry. Specifically, we introduced cysteine mutations in the RBD sequence in positions 68 and 172 of the NTD-RBD and RBD constructs to monitor conformations of the RBD. In contrast, we introduced cysteine residues in positions 1 and 68 of the NTD-RBD construct to monitor conformations of the NTD (**Fig. 1**). We will refer to these constructs as RBD_L_, NTD-RBD_L_, and NTD_L_-RBD respectively, where the L subscript identifies the region probed by the labels. All constructs have been expressed in *E.coli*, purified, and labeled with Alexa Fluor 488 and Alexa Fluor 594.

### Folding stability of RBD

As a preliminary step, we tested whether truncation of the NTD impacts the conformations adopted by the RBD and its folding stability, since this would alter the ability of the domain to interact with nucleic acids. Our previous single-molecule experiments^[7]^ showed that the RBD is equally stable when it is part of the full-length protein or of the isolated NTD-RBD construct, suggesting that the linker region does not impact its folding stability. Following this earlier work, we next directly measure the stability of the RBD in the absence of the NTD.

Single-molecule FRET measurements of the RBD construct show a single peak with high transfer efficiency (**Fig. 2**) that is compatible with previous observations of the completely folded RBD in the context of the NTD-RBD and full-length protein^[7]^. To confirm the observation, we further quantified the folding stability of the RBD in the absence of the NTD by titrating Guanidinium Chloride (GdmCl) into the RBD_L_ construct. Increasing the concentration of denaturant revealed the appearance of up to two species, which mirrors previous observations of an intermediate and unfolded state identified for the same domain^[7]^. An estimate of the relative abundance of each species can be computed by comparing the relative areas of the distinct populations. The data can be well described assuming a thermodynamic equilibrium between three states with ΔG_UI_ = 2.8 ± 0.1 kcal mol^−1^ and c_UI,1/2_ = 1.26 ± 0.03 M and ΔG_IF_ = 7.6 ± 0.4 kcal mol^−1^ and c_IF,1/2_ = 1.21 ± 0.01 M (**Fig. 2 and Supplementary Information**). Overall, our observations confirm that RBD is completely folded under aqueous buffer conditions. Compared to the full-length protein, truncation of the tail slightly shifts the unfolding transition towards lower GdmCl concentrations, but does not significantly affect the fraction folded in the absence of denaturant (**Supplementary Table 5**).

**Figure 2.**
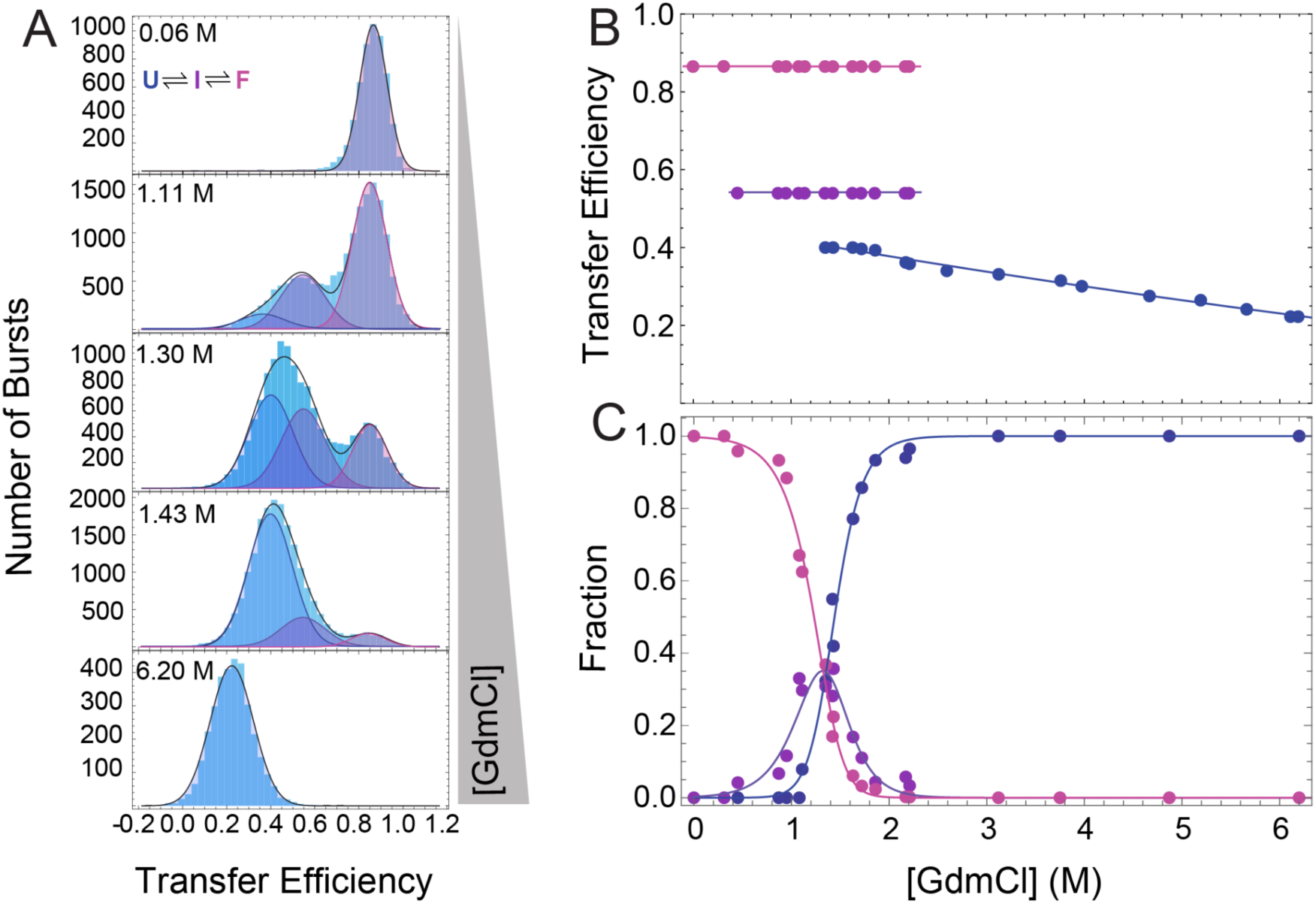
RNA Binding Domain (RBD) folding. **A.** Representative distributions of transfer efficiencies at different GdmCl concentrations. The transfer efficiency distributions are fitted with up to three Gaussian distributions. The folded configuration with high mean transfer efficiency is converted into an intermediate and unfolded state with lower mean transfer efficiencies with increasing GdmCl concentration. **B.** Mean transfer efficiencies obtained from a global fit of the histograms (see **Supplementary Information**) for the folded (magenta), intermediate (purple), and unfolded (blue) populations. Lines are guides for the eyes. **C.** Corresponding fractions of the folded (magenta), intermediate (purple), and unfolded (blue) populations. Lines represent a fit to the corresponding thermodynamic equilibrium according to **Eq. S3** and **S4.**

### Binding of nonspecific RNA to RBD

Given our goal is to quantify and compare the binding affinity of the RBD for RNA, we sought to develop a single-molecule assay that would let us quantify the fraction of bound protein as a function of RNA concentration. We first tested whether binding of RNA to RBD can be visualized via changes in transfer efficiency. With increasing concentration of a ∼200 nucleotide long poly(rU), we noticed a small but measurable shift toward higher values of transfer efficiencies, from a mean transfer efficiency of ∼ 0.87 to ∼ 0.90. (**Fig. 3**)

**Figure 3.**
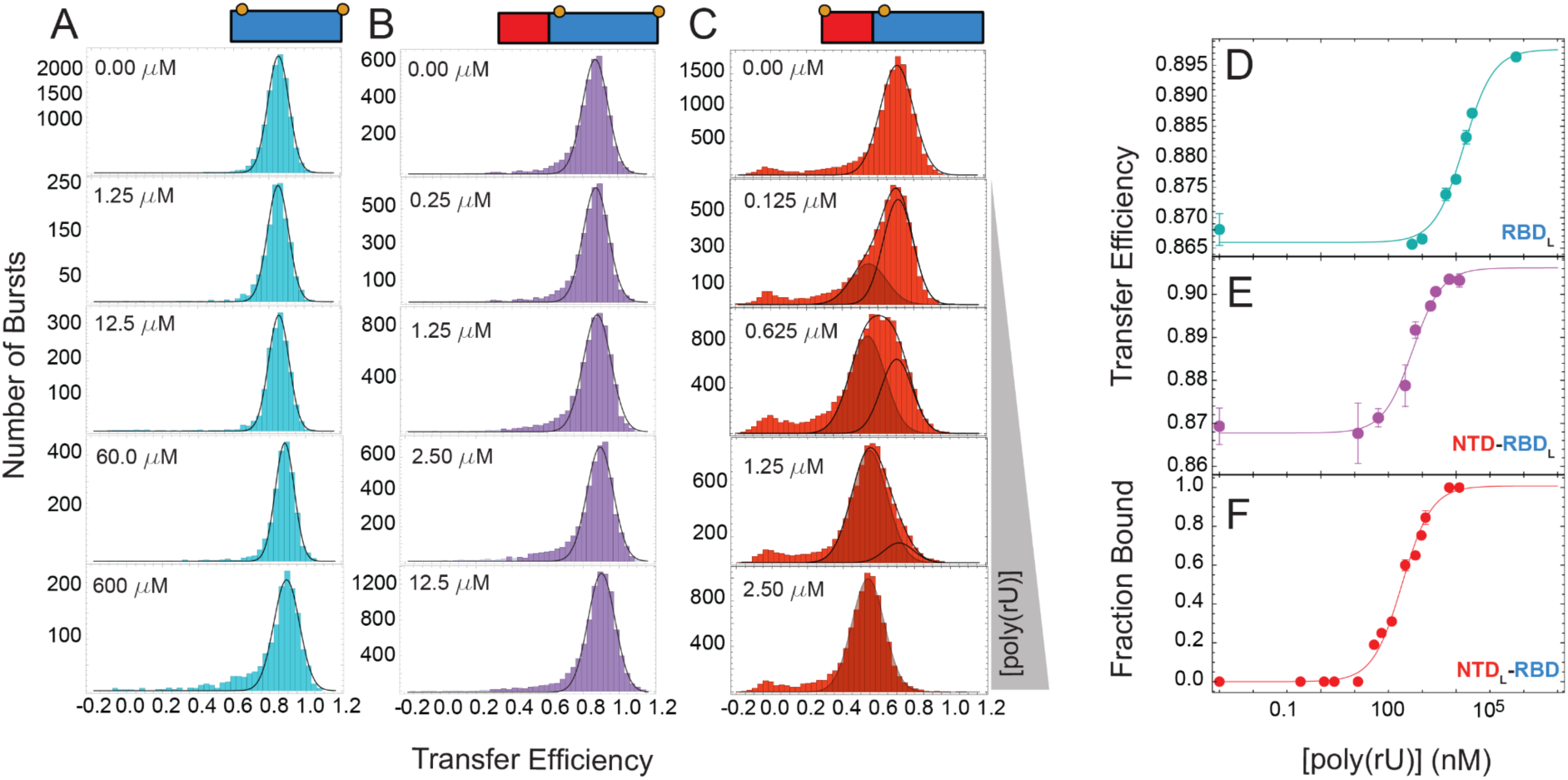
poly(rU) binding to RBD and NTD-RBD. **A.** Representative distributions of transfer efficiencies at different concentrations of poly(rU) for RBD_L_. Distributions are fitted to a single Gaussian distribution. **B.** Representative distributions of transfer efficiencies at different concentrations of poly(rU) for NTD-RBD_L_. Distributions are fitted to a single Gaussian distribution. **C.** Representative distributions of transfer efficiencies at different concentrations of poly(rU) for NTD_L_-RBD. Distributions are fitted to two Gaussian distributions. **D.**Variations in the mean transfer efficiency of RBD_L_ upon binding poly(rU). **E.** Variations in the mean transfer efficiency of NTD-RBD_L_ upon binding poly(rU). **F.** Fraction bound of NTD_L_ -RBD as a function of poly(rU) concentration. Solid lines represent the fit to the binding equations **Eq. 3**. Best fit values of *K*_int_ are shown in Supplementary Table 6.

A plot of the deviation in mean transfer efficiency as a function of nucleic acid concentration reveals a sigmoidal trend that saturates at high concentration, as expected for a binding isotherm of the RNA to RBD on a logarithmic scale. We note that in typical ensemble experiments, a 1:1 protein:nucleic acid binding stoichiometry cannot be automatically assumed when titrating a long nucleic acid with multiple binding sites against protein. However, here the 1:1 binding stoichiometry can be invoked because of the single-molecule nature of the experiments, where only labeled proteins are present in the solution and only one labeled protein per time is observed in the confocal volume. This is confirmed by Pulsed Interleaved Excitation, which provides a quantification of the labeling stoichiometry of the measured molecules and supports that the protein remains “monomeric” across the whole titration. This does not exclude the possibility of two unlabeled nucleic acids binding to the protein, though we would expect a change in the concentration-response (see for comparison binding of NTD-RBD_L_ to specific single-stranded RNA). A fit of the mean transfer efficiencies across the titration to the 1:1 binding model reveals an intrinsic association constant K_int_ of (6 ± 2) x 10^−2^ *μ*M^−1^ (**Fig. 3, Supplementary Table 6**) at the standard buffer conditions of 50 mM Tris, pH 7.4.

To further test whether the signal does indeed report on binding, we investigated the effect of nucleic acid length on the detected binding affinity. A decrease in the length of the nucleic acid is expected to result in weaker binding affinities because of the reduction in productive binding configurations for short oligonucleotides. When repeating the same titration, for (rU)_n_ oligonucleotides with length *n* = 10, 12, 15, 17, 20, 25, 30, and 40 nucleotides, we observe an analogous response of the transfer efficiency distribution, with the mean transfer efficiency increasing with increasing RNA concentration (**Fig. 4** and **Supplementary Fig. 1**). As for poly(rU), each titration curve can be well described by a 1:1 binding model and the corresponding equilibrium binding constants can be estimated. When plotted against the length of the oligonucleotide, a clear increase in the association constant *K_A_*(per molecule) is observed with increasing length of the RNA, ranging from (4 ± 3) x 10^−2^ µM^−1^ to (1.2 ± 0.3) µM^−1^ (**Supplementary Table 7**).

**Figure 4.**
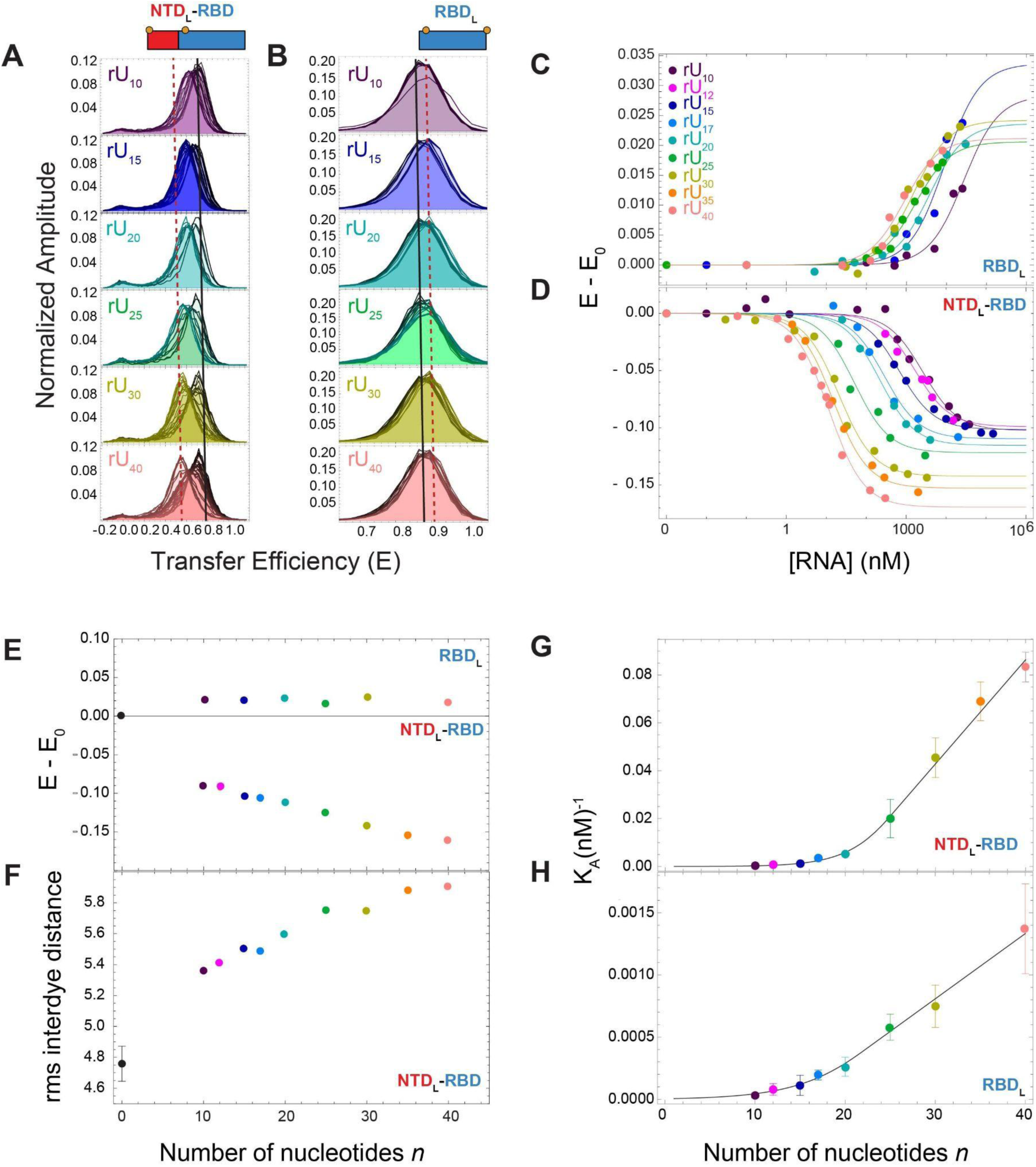
Length dependence of poly(rU) binding to NTD-RBD and RBD. **A-B.** Representative histograms of NTD_L_-RBD (**A**) and RBD_L_ (**B**) for rU*_n_* with nucleotide length *n* equal to 10, 15, 20, 25, 30, 40. The line of the transfer efficiency distribution varies from black (no RNA, starting condition) to the representative color of the specific length with increasing concentration of RNA. Black solid vertical line identifies the mean transfer efficiency at the starting condition (E_0_), red vertical dashed line identifies the mean transfer efficiency at “saturation”. **C-D.** Transfer efficiency changes upon (rU)_n_ binding, E-E_0_, for RBD_L_ (**C**) and NTD_L_-RBD (**D**) for all nucleotide lengths. Compare with single titrations in **Supplementary Fig. 1** for replicates and errors associated with each point. Solid lines are fit to **Eq. 1**. **E.** Variation range of transfer efficiency E with respect to the transfer efficiency E_0_ measured in absence of ligands for both NTD_L_-RBD and RBD_L_ constructs. **F.** Root-mean-square (rms) interdye distance of the disordered tail as measured by the labeling positions in NTD_L_-RBD and as a function of nucleic acid length. **G-H**. Association constants as a function of the number of nucleotide bases in (rU)_n_.

Assuming a simple unidimensional lattice model with an intrinsic association constant *K_in_*_t_, a given length of the nucleic acid *n*, and a contact site size of *M* nucleotides (the number of contiguous nucleotides involved in the interaction when a “complete” contact is realized with protein), we expect a linear trend as a function of *n* extrapolating through the x-axis (the length of the nucleic acid) at (*M* − 1), i.e.

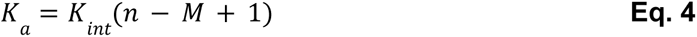

Indeed, measured association constants follow a linear trend and fit to **Eq. 4** results in an intrinsic association constant *K_int_* = (4.5 ± 0.5) x 10^−2^ µM^−1^ and a contact site size *M =* 12 ± 2. The model can be further developed to incorporate the contribution of partial interactions of the protein with the nucleic acid and include overhang effects, which in a first approximation can be described by:

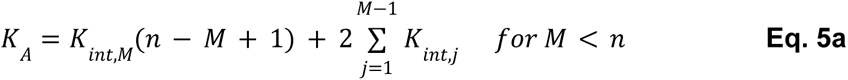

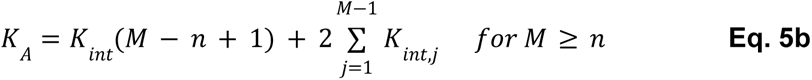

where *K_int,j_* represents a modified *K_int_*to account for the overhang effects (**Supplementary Information**).

The equation provides a quantitative representation of the complete dataset and identifies a *K_int_*= (5.2 ± 0.4) x 10^−2^ µM^−1^ and a contact site size *M =* 23 ± 2. Note that *K_int_* is within error of the value determined with **Eq. 4** and is consistent with the corresponding intrinsic association constant measured with the ∼200 nucleotide-long poly(rU). However, introducing partial binding at the ends of the chain leads to an increase in the estimate of the site size. This is a reflection of a strong assumption in the model, i.e. that the same average interaction is realized through all amino acids and nucleotides across the contact site (**Supplementary Information**). This obviously is an oversimplification that does not account for the contribution of ion release to the association constant as well as sequence-specific effects of the contact site. Therefore, the absolute value of the contact site size is likely to be overestimated by the fit to **Eq. 5**. The value falls between the estimates obtained with **Eq. 4** and **Eq. 5**. Having estimated the association constant and contact site size for the RBD, we then proceeded to investigate how the addition of the NTD alters these interaction parameters.

### Binding of nonspecific RNA to NTD-RBD

To test whether the addition of the disordered tail leads to a change in the binding affinity, we measured the association of the same poly(rU) using the construct NTD-RBD_L_. Titration of the RNA reveals a shift in the mean transfer efficiency that is analogous to the one observed for the RBD_L_, but the transition associated with binding is now shifted to low nanomolar concentrations. Fit of the mean transfer efficiency with a 1:1 binding model reveals a *K*_int_ = (2.0 ± 0.4) µM^−1^.

To confirm that this effect is due to the disordered tail, we turn to a second construct, the NTD_L_-RBD with labels in positions 1 and 68, which has been shown previously to report on the configurations of the disordered N-terminal tail and is in good agreement with the results from atomistic Monte Carlo simulations^[7]^. In the absence of RNA, this NTD_L_-RBD construct in aqueous buffer conditions reports on one narrow distribution that reflects the fast averaging over the conformational ensemble of the disordered tail. We proceed by testing if the same construct can report on RNA binding. With increasing concentration of poly(rU), we observe a modulation of the transfer efficiency distribution with a shift toward lower transfer efficiencies, from a mean transfer efficiency E = 0.709 ± 0.009 in absence of RNA to E = 0.542 ± 0.003 in presence of 10 µM of poly(rU) (**Fig. 3**). This observation clearly supports that the disordered tail is directly affected by the binding of RNA.

Analogous to the case of NTD-RBD_L_ and RBD_L_, an estimate of the binding affinity can be obtained by plotting the mean transfer efficiency (as fitted by a Gaussian distribution) as a function of the RNA concentration. Such analysis can be interpreted in terms of a simple 1:1 binding model, resulting in a *K*_int_ = (3.7 ± 0.4) µM^−1^. By a careful inspection of the width of the distribution, a broadening is observed for intermediate concentrations of RNA, suggesting that the measured distribution is indeed the resulting average of an unbound and bound population. Under this assumption, data can be refitted using two Gaussian distributions and the corresponding areas can be used to infer the fraction bound and unbound (**Fig. 3**). These quantities can be further analyzed to extract binding affinity for the nucleic acid, *K*_int_ = (4.0 ± 0.3) µM^−1^, which is in very good agreement with the one obtained from the mean value of the distribution. Both estimates of intrinsic association constants for the NTD_L_-RBD constructs are in close agreement with the one obtained for NTD-RBD_L_, confirming both constructs report on the same RNA binding independent of the labeling position. Based on these observations, the affinity of the NTD-RBD constructs appears to be ∼40-80 times tighter than that of the RBD alone, pointing to a direct contribution of the disordered region in favoring RNA binding.

Since the tail unequivocally favors binding, the conformations of NTD_L_-RBD upon RNA binding represent direct interactions of the tail with RNA. This poses a further question of whether the conformational change of the NTD represents a specific structural rearrangement due to an intrinsic encoded bound conformation or whether the conformational change reflects a dynamic conformational ensemble for the NTD-RBD/RNA complex. In the first case scenario, we expect that altering the length of the homo-polynucleotide sequence would possibly result in a change of affinity, but would not alter the mean transfer efficiency. In the second case scenario, instead, we expect to observe a change in both affinity and mean transfer efficiency.

To test this hypothesis, we investigated the binding of (rU)_n_ oligonucleotides with *n* ranging from 10 to 40 nucleotides (**Fig. 4**). For all of the sequences we observe a continuous shift in the mean transfer efficiency, reflecting binding of the RNA. Significantly, the mean transfer efficiency corresponding to the bound state depends on the length of the nucleic acid. The dependence of the mean transfer efficiency with the length of nucleic acid suggests a saturation effect that is reached for sufficiently long RNA. Inspection of the binding equilibrium constant as a function of length reveals two distinct regimes, which - as a first approximation - can be described by using **Eq. 4** and **2.** A linear fit using **Eq. 4** for RNAs with length between 20 and 40 nucleotides results in a *K*_int_ = (4.2 ± 0.4) µM^−1^ and *M* = 21 ± 1 nucleotides. A complete fit of the dataset using **Eq. 5** results in an intrinsic association constant *K*_int_ = (4.3 ± 0.2) µM^−1^ and *M* = 25 ± 2 nucleotides. The change in slope at approximately 20 nucleotides indicates that this length of nucleic acid is required to satisfy all the contacts between the nucleic acid and the NTD_L_-RBD construct, which results in a larger contact site size. In addition to a larger contact size, the interaction per nucleotide is tighter than the one determined for the RBD alone, as indicated by the NTD-RBD K_int_. Interestingly, a shift in transfer efficiency is observed for lengths shorter than the contact site size of RBD, implying that even for short oligos not all the contacts occur within the folded domain, and interactions with the tail need to be formed.

Taken together with the tighter *K*_int_ observed for NTD-RBD, these observations indicate that the complex between RNA and NTD-RBD is not solely initiated by contacts with the RBD domain but instead relies on dynamic interactions between the RNA and both RBD and NTD. Furthermore, the transfer efficiency shift does not saturate at the contact site size of the NTD-RBD construct (20 nucleotides); instead, a continuous change is observed for longer lengths, approaching saturation at approximately 40 nucleotides. These observations further suggest a dynamic complex between the protein and RNA, where the position of the contacts formed depends on the number of available nucleotides and the contact site size represents a mean number of minimum contacts that are formed above a given length of the oligo.

To test this hypothesis, we performed ns-FCS measurements of the NTD_L_-RBD in the presence of RNA. We previously showed that the NTD region in absence of RNA is flexible and dynamic^[7]^. ns-FCS measurements of the NTD_L_-RBD in the absence of RNA reveals a reconfiguration time of approximately 110 ± 20 ns, which is marginally affected upon binding RNA, with a reconfiguration time of the NTD spanning a range between 94 and 108 ns across the different lengths tested from (rU)_10_ to (rU)_40_ (**Supplementary Fig. 2**). This indicates that the NTD remains largely dynamic and contacts must occur only across a small set of nucleotides.

### Simulations of RNA binding to NTD-RBD

To gain a molecular understanding of the interaction between RNA and the NTD-RBD, we turned to coarse-grained molecular dynamics simulations. We utilized the Mpipi force field, a recently-developed model that combines short-range interactions and long-range electrostatics and encodes each amino acid or nucleotide as a chemically-distinct entity (**Fig. 5A**) ^[28]^. Mpipi was specifically developed with intrinsically disordered regions in mind^[28]^. Previous work has shown good agreement between simulations and experiments when this model has been used to assess non-specific protein-protein and protein-RNA interactions leading to phase separation ^[28,33,34]^. We first simulated RBD with (rU)_10_ to identify residues on the folded domain that contribute to ssRNA binding (**Fig. 5B**). We calculated protein:RNA contacts from these simulations and observed reasonable agreement with previously-reported NMR chemical shift perturbation experiments of the RBD with ssRNA, performed with a 10-mer RNA of 5’-UCUCUAAACG-3’^[31]^. This result suggests that our simulations, at least qualitatively, are able to recapitulate experimentally measured protein:RNA interactions (**Fig. 5B**).

**Figure 5.**
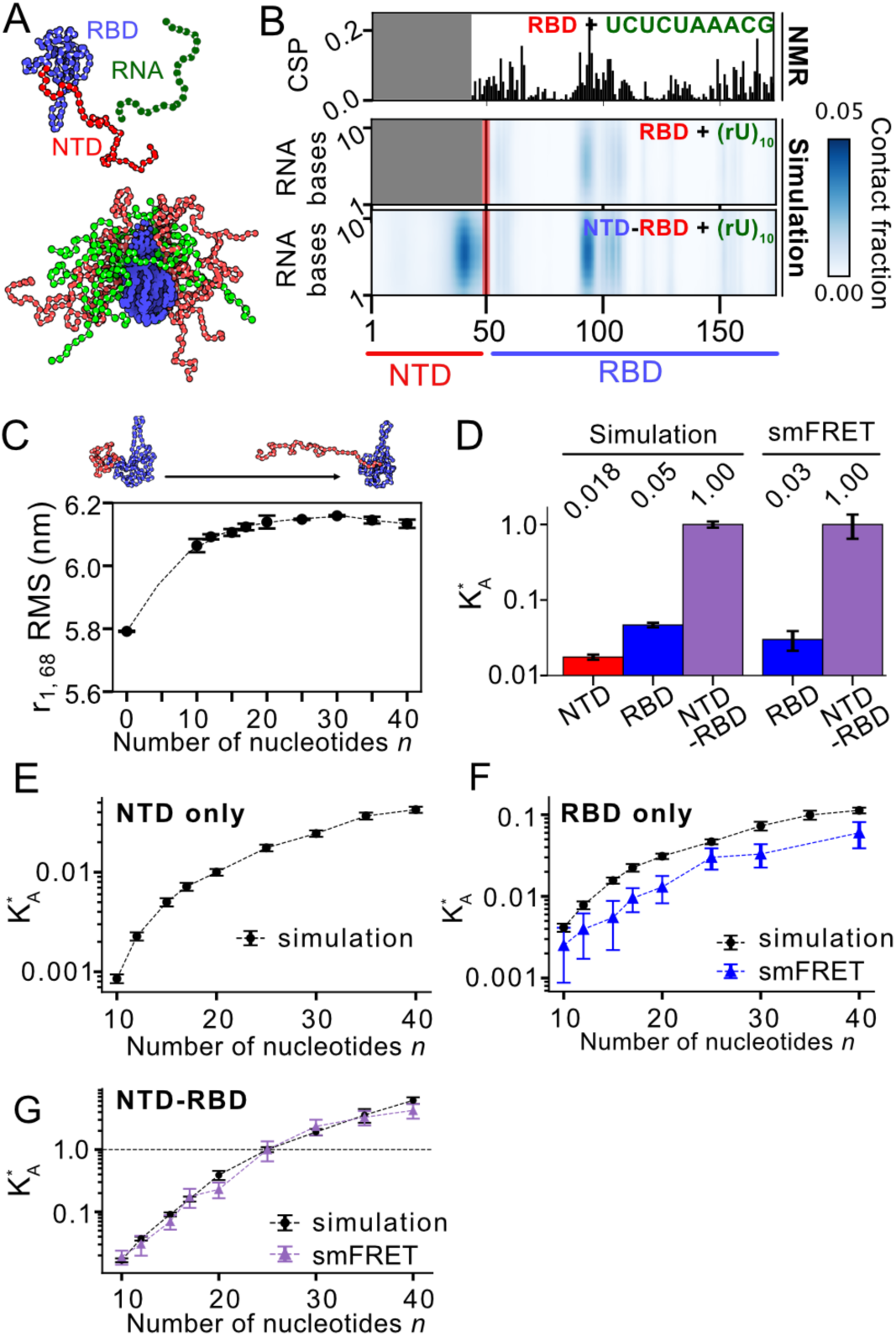
Coarse-grained simulations of the Nucleocapsid protein with ssRNA. **A.** The Mpipi forcefield is used to model SARS-CoV 2 N-protein interactions with ssRNA (rU) ^[28]^. Each amino acid and nucleotide is represented as a single bead (see Methods). The Nucleocapsid-RNA bound state is highly dynamic (bottom). **B.** Simulations of RBD + (rU)_10_ (middle) or NTD-RBD + (rU)_10_ (bottom) enable the assessment of which residues engage in direct RNA interactions. Protein:RNA contacts are quantified by calculating the contact fraction, defined as the fraction of the simulation in which each amino acid-nucleotide pair is under a threshold distance of 14 Å. The specific threshold chosen does not alter which residues are identified as RNA-interacting (**Supplementary Fig. 3**). The pattern of residues identified from simulations shows qualitative agreement with chemical shift perturbation data of the RBD (amino acids 44-173) observed upon binding to a 10-mer ssRNA (5’- UCUCUAAACG-3’)^[31]^. **C.** Root-mean-square distance (RMSD) between residues 1 and 68 increases upon ssRNA binding, with a modest increase observed in the RNA-bound state as a function of RNA length up to (rU)_20_. **D** The normalized binding affinity (K_A_*) of the NTD, RBD, or NTD-RBD binding to (rU)_n_ is calculated as the apparent binding affinity divided by the apparent binding affinity for NTD-RBD binding (rU)_25_. K_A_* can be calculated in a self-consistent manner for simulations (left) and experiment (right). **F** Length dependent K_A_* of the NTD + (rU)_n_. **G** Length dependent K_A_* of the RBD + (rU)_n_. **H.** Length-dependent K_A_* of the NTD-RBD + (rU)_n_. For **F**,**G** and **H,** K_A_* is calculated by dividing the apparent K_A_ from the specific (rU)_n_ length by the apparent K_A_ from the NTD-RBD + (rU)_25_ simulation.

Having first performed simulations of (rU)_10_ with the RBD, we next performed simulations of NTD-RBD and (rU)_10_. In addition to the previously observed RBD interactions with (rU)_10_, we now observed additional interactions between the disordered NTD and (rU)_10_ (**Fig. 5B, Supplementary Fig. 3**). The NTD remains fully disordered in the bound state of NTD-RBD:(rU)_10_ (**Supplementary Fig. 4**) and the pattern of RBD – (rU)_10_ interactions is comparable in both the presence and absence of the disordered NTD. While the same RBD residues engage with RNA in the presence vs. absence of the NTD, the frequency is altered. Specifically, the NTD enhances interactions between residues 89 – 107 of the RBD with RNA (**Supplementary Fig.3**). This region maps to the β-extension previously identified as engaging in RNA interactions ^[31]^. Within the NTD, residues 30-50 contain five positively charged amino acids (four arginines and one lysine) and interact directly with (rU)_10_, in good agreement with recently published NMR experiments^[35]^ (**Fig. 5B**). Taken together, these results suggest that the presence of the NTD potentiates RBD:RNA interactions as well as engaging in a new set of interactions with RNA.

We then tested whether our simulations capture the enhanced affinity of NTD-RBD with RBD and the length dependence of the binding model. By defining the fraction of the simulation in which the protein and RNA are bound to one another, we can calculate an apparent binding association constant (K_A_) for simulations with either RBD or NTD-RBD and compare the relative values (see **Supplementary Tables 8-9** and **Supplementary Fig. 5**). Comparing the binding of these two constructs to (rU)_25_ (which is larger than the measured contact site size of RBD and equivalent to the upper limit of the one of NTD-RBD), the presence of the NTD increases the K_A_ by a factor of 4.7 ± 0.4 %, in good agreement with our experimentally measured ratio of association constants of 3 ± 1 % for K_A,RBD_/K_A,NTD-RBD_ (**Fig. 5D, Supplementary Table 10**). Intriguingly, simulations of NTD alone with (rU)_25_ revealed substantially weaker binding compared to either the RBD or NTD-RBD (**Fig. 5D-G**). This suggests that the NTD’s ability to enhance RNA binding – at least in the context of poly(rU) – is an emergent property of the NTDs location relative to the RBD, as opposed to solely an intrinsic ability to bind RNA tightly.

Our single-molecule FRET experiments revealed an expansion of the NTD upon binding to RNA, where longer single-stranded RNAs lead to a higher degree of NTD expansion (**Fig. 4**). This is in contrast to simple expectations for polyelectrolyte condensation, where oppositely charged polymers are expected to compact upon interaction with one another^[36,37]^. This RNA-dependent expansion of the NTD is reproduced in our simulations, where we observed an increase in the root mean square distance (RMSD) between residues 1 and 68 of the NTD upon RNA binding, followed by a modest increase in RMSD as the RNA length increases up to (rU)_20_ (**Fig. 5C**). These trends are in qualitative agreement with the single-molecule FRET measurements (**Fig. 4F**). These results confirm our ability to capture the conformational behavior of the NTD upon RNA binding, while adding further evidence of RNA length dependent expansion of the NTD.

Importantly, in all the simulations the bound state is a dynamic complex that is compatible with the dynamics observed in nsFCS experiments (**Fig. 5A, Supplementary Movie 1**). Taken together, our results suggest that NTD-RBD interacts with RNA forming a disordered “fuzzy” complex largely driven by the interaction with positively charged groups in the NTD and RBD.

### Effect of salt

Protein-RNA interactions are known to be sensitive to salt concentrations due to the large contribution of electrostatics. A significant contribution to binding can arise from condensed ions on protein and RNA, which can be released upon binding. To estimate the extent of ion release, we measured the association constant as a function of the salt concentration. We restrict our investigation to (rU)_20_ and (rU)_40_, where we can quantify affinities up to 200 mM KCl in the range of available concentrations of the ligand. As shown in **Supplementary Fig. 6 and 7**, the mean transfer efficiency of the NTD_L_-RBD is marginally altered by salt screening in absence of the ligand, which is consistent with previous observations^[7]^.

NTD_L_-RBD was titrated with (rU)_n_ at different KCl concentrations. Representative histograms and the observed dependence of *K*_A_ on salt concentration are shown in **Fig. 6** and **Supplementary Fig. 6-7**. Both (rU)_20_ and (rU)_40_ datasets reveal a linear trend on the log-log plot of *K*_A_ and K^+^ concentration. Analogous results are obtained when considering the total concentration of cations K^+^ and Tris^+^ (**Supplementary Fig. 8**). The lack of curvature in K^+^ titration suggests that interactions with Tris^+^ ions do not contribute substantially to ion release. The slope of the linear trend is equal to -5.1 ± 0.4 and -5.0 ± 0.5 for (rU)_20_ and (rU)_40_, respectively, indicating a net release of ∼ 5 ions upon interaction^[38]^ (see **Supplementary Table 11**). Finally, our measurements also provide a quantification of the RNA binding association constants at the physiological concentrations found in cells (∼150 mM K^+^). When compared to corresponding values observed in the reference buffer condition, we observe an decrease of the association constant *K*_A_ to (0.17 ± 0.02) µM^−1^ for (rU)_20_ and (0.38 ± 0.04) µM^−1^ for (rU)_40_, corresponding to a weaker affinity in higher salt concentration (see **Supplementary Table 12**).

**Figure 6.**
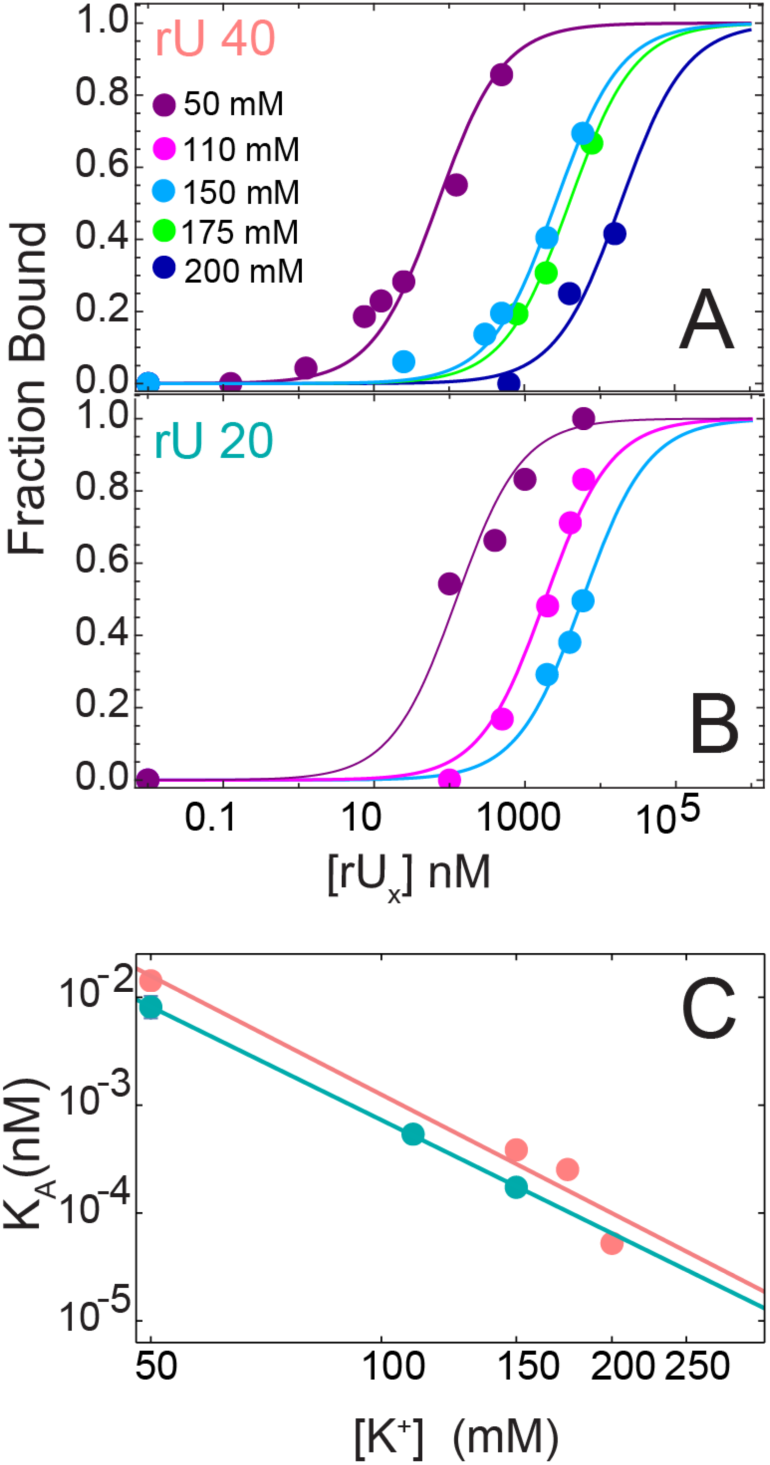
Salt dependence of binding association constant. Fraction bound is determined from single-molecule FRET experiments of the NTD_L_-RBD as a function of (rU)_40_ (**A**) and (rU)_20_ (**B**) concentration. Each curve is measured in 50 mM Tris buffer and increasing KCl concentration: 50 mM (purple), 110 mM (magenta), 150 mM (cyan), 175 mM (green), 200 mM (blue) KCl. See corresponding histograms in **Supplementary Fig. 6-7** and **3**. Solid lines are fit to **Eq. 2a**. **C.** Association constants determined from the measurements in panel A ((rU)_40_, pink) and panel B ((rU)_20_, cyan) are plotted against the concentration of K^+^ ions on a log-log plot. Solid lines represent the linear fit of Log(*K*_A_) as a function of Log([K^+^]). Results for total ion concentration are reported in **Supplementary Fig. 8**. The similar slope of (rU)_40_ and (rU)_20_ data suggests that the same net ion release occurs upon binding of the two different lengths of nucleic acids (see **Supplementary Table 11**).

### Interaction with specific single-stranded RNA

To test whether sequence specificity can affect affinity and mode of binding of the specific RNA with the disordered region, we studied the interactions with a 21 nucleotide sequence (V21) from the 5’ UTR of the viral genome. This region of the genome was previously found interacting with the N protein in *in cell* crosslinking studies ^[10]^ and has been confirmed to adopt no secondary structure at room temperature^[39]^.

We quantified binding of V21 using the NTD_L_-RBD construct. As for the case of nonspecific single-stranded RNA, at increasing concentration of V21, we notice a shift of the mean transfer efficiency that reaches a saturating value at ∼ 1 µM RNA concentration, which we interpret as representing the binding between one protein and one RNA strand. However, at concentrations of V21 higher than 1 µM, we observe the appearance of a second population at lower transfer efficiency, which is consistent with a second binding event of the nucleic acid to the protein, i.e. a 2:1 RNA:protein stoichiometry. This conformational change is associated with a mean transfer efficiency that is significantly lower than any of the mean transfer efficiencies that has been observed for poly(rU) (E ∼ 0.37), indicating a distinct mode of binding and structural organization of the NTD. We interpret such an extended configuration as an expansion of the tail to accommodate two nucleic acid molecules. Since we observe this second mode of binding only for V21 but for none of the poly(rU) sequences, we propose that this second bound state is the result of a partial hybridization of the V21 sequence.

To quantify the association constants corresponding to the different binding events, we globally fit the change in the mean transfer efficiency associated with the first binding event and the change in relative area of the second population associated with the second binding event (**Fig. 7, Supplementary Table 13**). Data are globally fit to a model that accounts for two distinct bound states with corresponding association constants *K* ^V21^ of (6.2 ± 0.3) µM^−1^ and *K* ^V21^ of (0.15 ± 0.10) µM^−1^. *K* ^V21^ is ∼ 50% larger than the corresponding association constant for r(U)_20_, *K* ^rU20^ = (4.3 ± 0.3) µM^−1^, whereas the mean transfer efficiency of the bound state appears only slightly smaller than that for r(U)_20_. To better understand if the second mode of binding is compatible with double-stranded sequences, we turned to the investigation of specific double-stranded RNA sequences.

**Figure 7.**
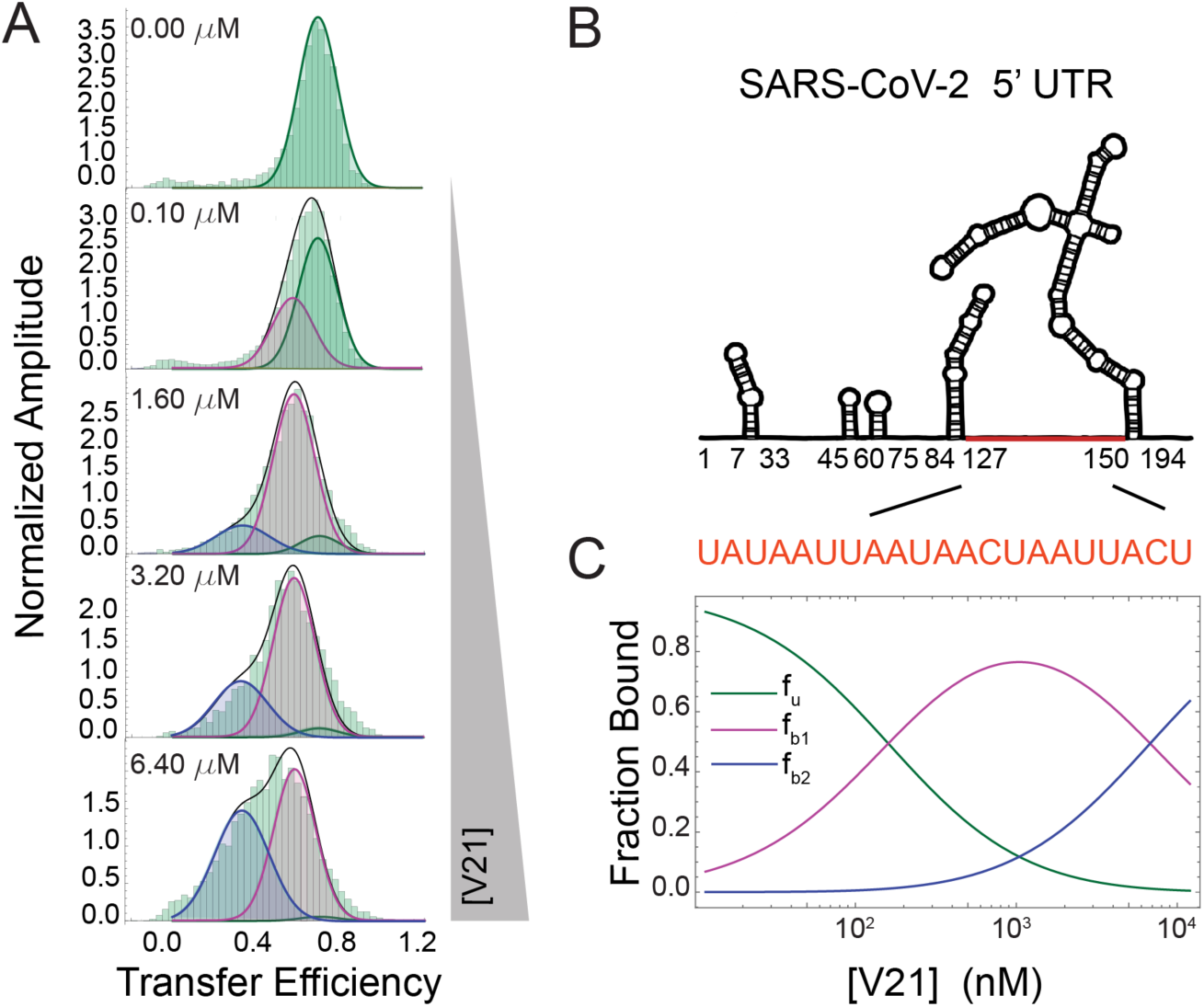
Specific ssRNA binding to NTD_L_-RBD. **A.** Representative distributions of transfer efficiencies upon binding of V21. Increasing concentration of RNA leads to a first conformational change of the tail that appears to be largely completed at ∼3 µM. Further increasing the concentration of V21 leads to a second conformational change of the disordered region, indicating that the protein is binding two copies of the nucleic acids. Areas are fitted according to **Eq. 2b** and **2c**. **B.** Graphical representation of the SARS-CoV-2 5’ UTR based on Iserman et al.^[10]^, highlighting the region corresponding to V21. **C.** Fraction of each state: unbound (f_u_), bound to one V21 molecule (f_b1_), and bound to two V21 molecules (f_b2_). Corresponding values of the fit are reported in **Supplementary Table 13**.

### Interaction with specific RNA hairpins

The 5’ UTR of the SARS-CoV2 genome contains short single-stranded regions and various conserved hairpins, which can offer additional binding sites to the NTD-RBD. In addition, double-stranded regions of the genomic RNA have been proposed as putative packaging signals^[27]^, including the SL5B hairpin in the 5’ UTR and the NSP15 hairpin from the mRNA of the Nonstructural Protein 15^[27,40,41]^ (see **Fig. 8, Supplementary Fig. 9, Supplementary Table 14**). Given the potential role of these regions in driving condensation of the nucleic acid, we focused on these two archetypal sequences. NSP15 and SL5B were transcribed *in vitro*, and their hairpin structure at room temperature was confirmed by thermal melting experiments (**Supplementary Fig. 10**).

**Figure 8.**
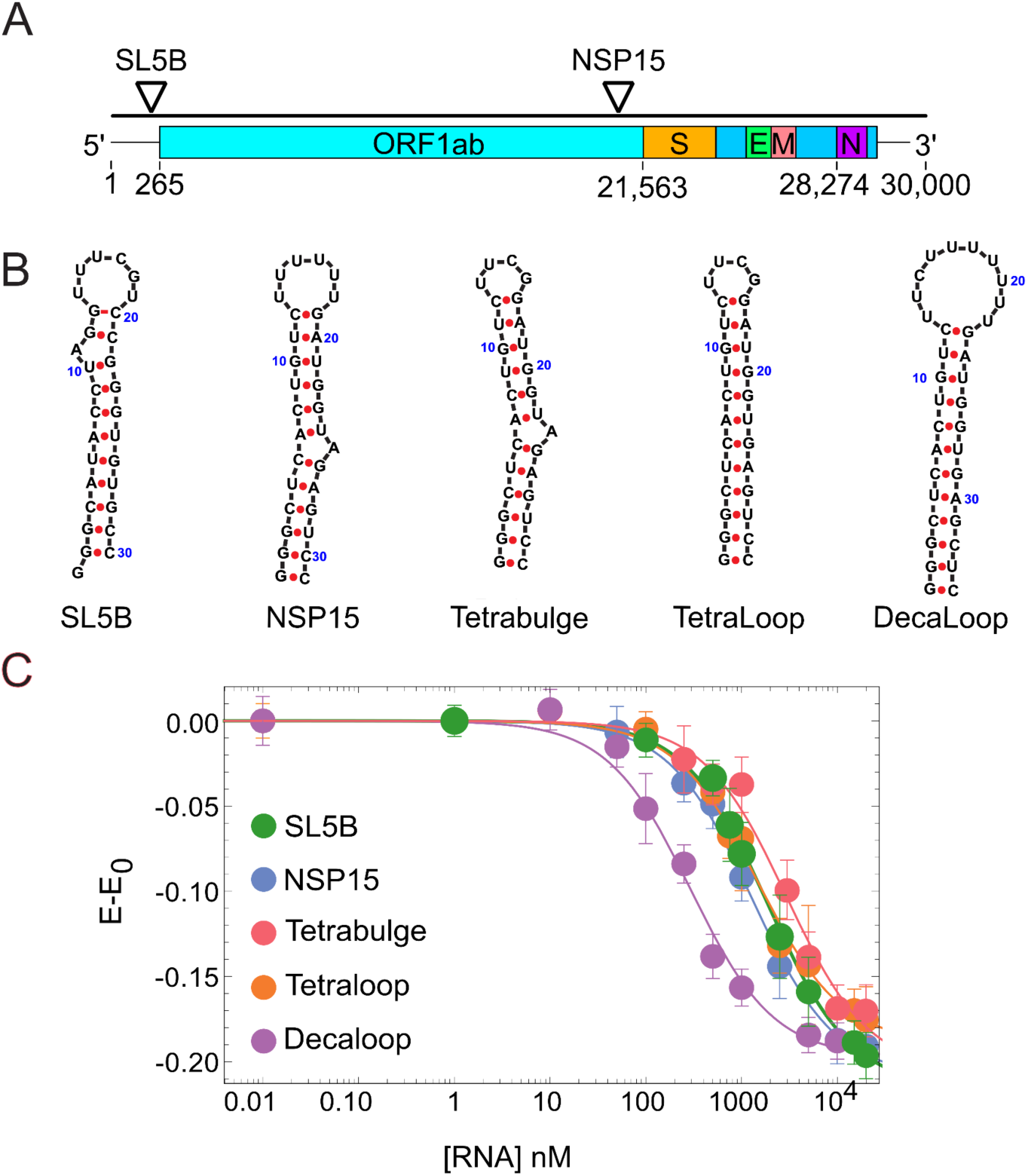
Specific hairpin RNA (hpRNA) binding to NTD-RBD. **A.** Position of studied hpRNA sequences in the viral genome. **B.** Hairpin structure and sequence. **C.** Variation in the mean transfer efficiencies of the NTD_L_-RBD as a function of hpRNA concentration. When no hpRNA is present, transfer efficiency is ∼0.68 (compare with **Supplementary Figure 8**). Solid lines are fit to **Eq. 1**.

Single-molecule FRET measurements of the NTD_L_-RBD construct bound to either SL5B or NSP15 reveal a clear shift of the transfer efficiency distribution toward lower values, i.e. more extended configurations. Deviation of mean transfer efficiency can be fit as in the case of single-stranded RNA to determine the association constants: *K* ^NSP15^ = (7.8 ± 0.7) x 10^−1^ µM^−1^ and *K* ^SL5B^ = (5.3 ± 0.4) x 10^−1^ µM^−1^. These values are compatible with the one associated with the second binding mode of V21, *K* ^V21^, supporting the hypothesis that this binding mode is due to hybridization of a double-stranded RNA. Interestingly, the conformational changes of NTD_L_-RBD bound to the hairpins appear to be larger than what is observed for the majority of single-stranded RNA, even if the binding affinity is weaker. We attribute the increased expansions of the disordered tail to the larger excluded volume of the double-stranded hairpin.

Finally, we turned to investigate which regions of the hairpins may contribute to the binding. Due to the similar affinity of these sequences to that of (rU)_10_, we hypothesized that NTD-RBD may preferentially bind to the RNA hairpin through its loop region. We chose the NSP15 sequence as a reference and designed RNA hairpins (hpRNA) with perfect duplex stems and loops of either 4 or 10 nucleotides (**Fig. 7**). We refer to these constructs as TetraLoop and DecaLoop. The four nucleotide loop in the TetraLoop is cUUCGg, and is expected to result in a unique and stable structure, while the ten nucleotide loop contains seven U’s and is unlikely to form internal structure. We found that the binding affinity of these two hpRNAs does seem to depend on the length of the loop, with a *K* ^Tetraloop^ = (6.7 ± 0.8) x 10^−1^ µM^−1^ and a *K* ^Decaloop^ = (3.4 ± 0.5) µM^−1^, suggesting that the single-stranded loop does influence the affinity and, therefore, could be the main site of interaction. However, affinity is stronger than that of (rU)_10_, indicating that binding involves both single- and double-stranded regions of the nucleic acid.

To probe the possible roles of defects in double-stranded regions, we tested whether introducing an unpaired A in the tetraloop hairpin stem would affect binding. We do not find significant differences from the perfect stem (*K*_A_^Tetrabulge^ = (3.4 ± 0.7) x 10^−1^ µM^−1^), suggesting that small defects in the duplex do not influence the NTD-RBD region. Larger internal loops could act as binding sites, but these would depend on sequence and context.

### Omicron variant

Many mutations in the N protein occur within the disordered regions^[42]^. The Omicron variant offers a convenient point of comparison, with three key mutations found in the NTD. More than 90% of sequences on the GISAID database (accessed on February 8 2023) report a proline to leucine substitution in position 13 and deletion of three residues between positions 31 and 33^[43]^ (**Supplementary Table 2**). Residue 13 is part of a predicted short helix motif ^[7]^ that may offer an interaction site for RNA binding, whereas residues 31 and 32 contain two oppositely charged residues. To test the impact of these mutations, we expressed, purified, and labeled the Omicron NTD_L_-RBD (^Om^NTD_L_-RBD).

We first characterized the conformations of the tail in absence of RNA. Given the small variations in the sequence, both in terms of hydrophobicity and net charge, we expect negligible variations. Indeed, we observed no significant shift in transfer efficiency (**Fig. 9**). We then performed binding experiments at increasing concentrations of poly(rU). We observed an identical mean transfer efficiency at saturation concentrations of poly(rU) and *K*_A_ = (9 ± 1) 10^−1^ µM^−1^, approximately 4 times weaker binding affinity than for the wild-type sequence. These observations overall support that the mode of binding of RNA is similar between NTD_L_-RBD (Wuhan-Hu-1) and ^Om^NTD_L_-RBD (as supported by the same transfer efficiency in the bound state), but with different affinities (as indicated by the concentration dependence).

**Figure 9.**
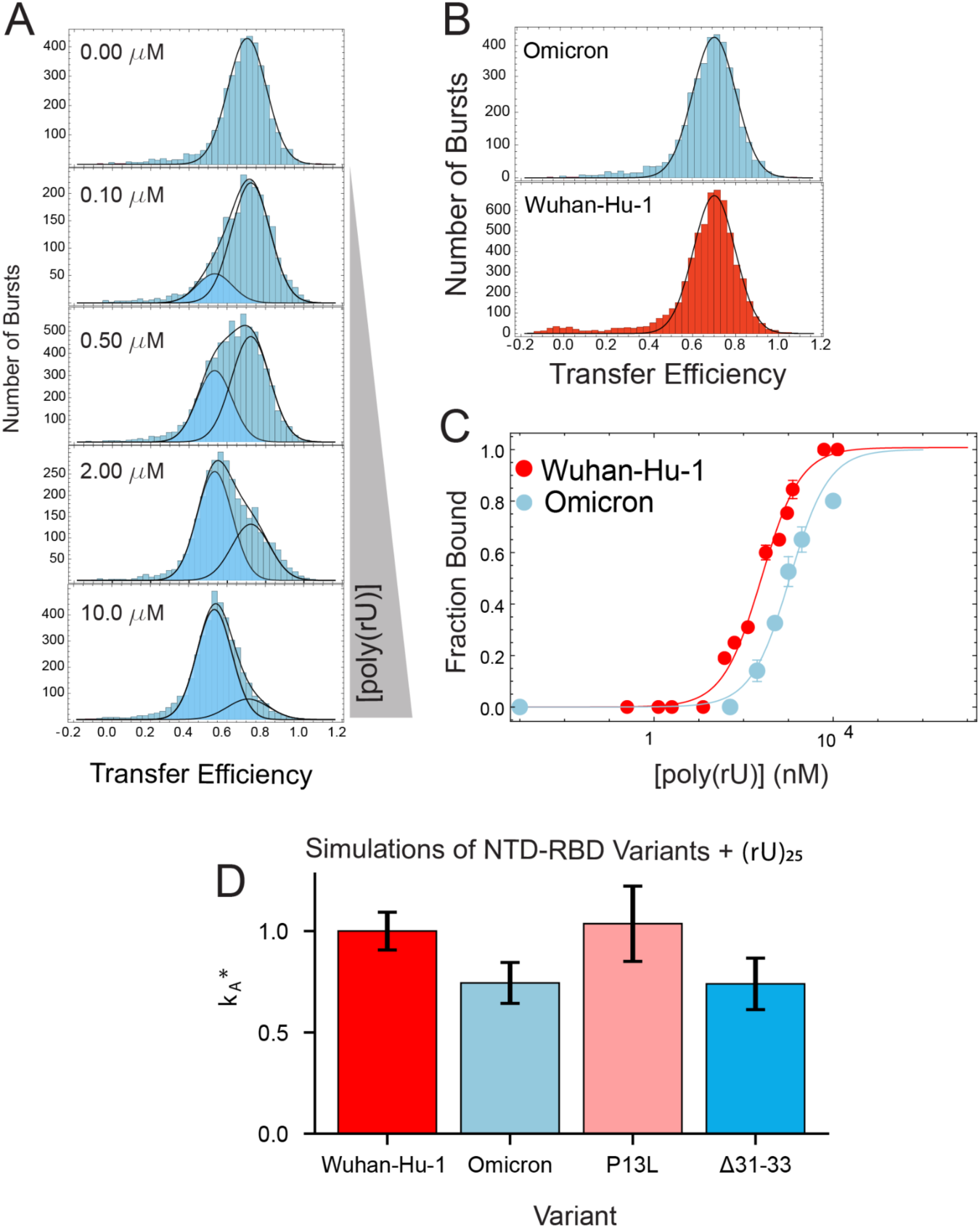
Omicron variant. **A.** Transfer efficiency distributions for the Omicron variant as function of poly(rU) concentration. Distributions are fitted with up to two Gaussian distributions to quantify the mean transfer efficiency and relative fraction of bound and unbound fractions. **B.** Comparison of unbound configuration of disordered tail for Wuhan-Hu-1 (red) and Omicron variant (cyan) reveals no significant variations in overall conformations. **C.** Comparison of binding affinity for Wuhan-Hu-1 (red) and Omicron variant (cyan) reveals different affinities for poly(rU). Solid lines are fit to **Eq. 2a**. **D.** Trend of the normalized binding affinity (*K*_A_*) predicted by simulations with Mpipi model for the Omicron mutant and additional variants.

We further investigate molecular insights by performing corresponding coarse-grained simulations. Here, we observed a decrease in binding affinity between Wuhan-Hu-1 and the Omicron variants. We then tested whether this difference is driven by the lack of the proline substitution or by the charge suppression (**Fig. 9**). Mutating only the proline to leucine in our simulations resulted in no detectable change in the binding affinity. In contrast, maintaining the proline and deleting residues 31 to 33 results in a suppression of binding affinity, suggesting that the change in RNA binding affinity observed for Omicron NTD-RBD is dominated by charge effects **(Supplementary Table 15)**. Overall, our observations indicate that small changes in the sequence composition of NTD may not alter the overall conformational behavior of the chain, but can significantly impact the binding affinity.

## DISCUSSION

### The NTD is essential for RBD function

The N protein is responsible for packaging the SARS-Cov-2 genome, but the molecular mechanism of this process remains underdetermined. While previous work has focused on folded domains of the protein as possible centers for interactions, here we have been exploring the role played by one of the disordered regions to determine if the disordered region is a disposable appendage to the folded domain or plays a role in determining protein function. In particular, we investigated the NTD-RBD region and quantified how the disordered NTD contributes to the mode of binding and affinities for RNA. Through our experiments, we have discovered that the RBD alone binds very weakly to single-stranded RNAs, while the NTD significantly increases RNA binding affinity. Altogether, our data suggest that the RBD alone cannot be considered a primary determinant of RNA binding, and association is most likely the result of the concerted interaction of the RBD and surrounding disordered regions with RNA.

### The NTD-RBD forms a dynamic complex with RNA

Our data confirm the previous observations that the NTD is a flexible and dynamic region^[7]^, whose large degree of conformational heterogeneity is retained when the protein is bound to RNA. Thus in defining the interactions between the NTD and RNA, we cannot model the complex as a rigid body with fixed interactions; rather, we have to consider the points of interaction that can be sampled by the disordered protein and nucleic acid. Inspection of the sequence composition (**Supplementary Table 1**) reveals 7 positive charged residues (6 Arg and 1 Lys) and 2 hydrophobic residues (1 Phe and 1 Trp), which offer possible sites of interaction with the nucleic acid. Indeed, arginines can neutralize phosphate groups on the RNA and aromatic groups of Phe and Trp can stack with RNA bases. From a point of view of the sequence pattern, two Arg and one Phe residues occur in a putative helix (identified in our previous simulations^[7]^) that span from residue 10 to 16, one Trp and Phe are positioned at the junction between the NTD and RBD, and the remaining Arg and Lys residues are clustered between position 30 and 50.

Our coarse-grained simulations point to a key role of electrostatic interactions in regulating the binding of the nucleic acid to the NTD-RBD region, in particular, the stretch between residues 30 and 50 in the NTD and between residues 85 and 110 in the RBD (**Supplementary Fig. 11 and 12**). These RBD residues comprise the positively charged β-extension, a flexible pair of beta strands that prior work has identified as wrapping around single-stranded RNA during binding ^[31]^. Previous computational work proposed that the interplay between charged residues on the RBD surface and in the NTD can tune NTD conformational behavior^[44]^. An additional explanation for these previous observations could be one in which N protein has evolved across coronaviridae to ensure high-affinity RNA binding, with compensatory/co-evolutionary changes in the NTD and RBD ensuring that non-specific electrostatically-driven interactions are conserved in spite of sequence variation in both the NTD and RBD.

Our simulations also allow us to deconvolve the relative contributions of the NTD and RBD to RNA binding, illustrating the benefit of a combined, multi-pronged approach in molecular dissection ^[45]^. Although the addition of the NTD to the RBD leads to a substantial increase in binding affinity, our simulations predict that, in isolation, the NTD binds RNA more weakly than either the RBD or the NTD-RBD. With this in mind, the impact of the NTD appears to be mediated by its position relative to the positively-charged β-extension on the RBD. The resulting orientation offers a dynamic, positively charged binding surface, such that the emergent binding affinity is substantially higher than would be naively expected, likely through both an avidity effect and by prepaying the entropic cost of bringing two positively charged protein regions into relatively close contact with one another.

In addition, the simulations corroborate the experimental intuition of a dynamic complex where not only the protein but also the nucleic acid is exploring heterogeneous conformations in the bound state. Overall, these observations ascribe the NTD-RBD:RNA complex to the category of so-called “fuzzy” complexes. The strong electrostatic nature of the interactions is consistent with the recent observation of highly dynamic complexes formed by oppositely charged biopolymers^[46]^, as for the case of prothymosin alpha and histone H1^[47,48]^.

### The NTD-RBD region prefers single-stranded RNA

Our data clearly support the conclusion that the NTD-RBD exhibits some discrimination among RNA targets. We find a generally higher affinity for both specific and non-specific sequences of single-stranded RNA. This is consistent with previous studies of N protein^[10,49]^, including *in cell* crosslinked studies of the protein to the 5’ UTR^[10]^, where single-stranded regions, several large loops and junctions predominated the interactions. Additional studies also identified short U-tracts as possible targets of the interaction. Compared to single-stranded RNA, our work finds lower affinities for double-stranded RNA sequences. In particular, our investigation of model hairpins based on the NSP15 genome region tested the role of RNA duplexes, hairpin loops, and duplex deformations in NTD-RBD association. We found that small deformations in the duplex do not significantly alter the interaction with the protein, whereas an increase in the size of the loop region results in an increase of the binding affinity, confirming a preferential interaction of this protein region with single-stranded RNA.

### NTD mutations alter RNA binding

A high number of mutations occur in disordered regions of the Nucleocapsid protein^[42]^. Our results on the impact of the Omicron NTD mutations clearly show that alterations of three amino acids in this IDR are sufficient to decrease the interaction affinity between the construct and the nucleic acid. This implies not only that the N protein IDRs play a role in the interaction of the protein with nucleic acids, but that mutations in the same regions can effectively alter the function of the protein. Moreover, while it is often assumed that small changes in IDRs may not substantially influence molecular function, our results here provide a clear counter-example, whereby a 4-times change in binding affinity is driven by just a few mutations. The sensitivity of RNA binding to small sequence changes that alter the charge of the protein also raises the possibility that phosphorylation may play a role in tuning RNA binding affinity, as has been proposed previously ^[8,50]^.

The fact that mutations minimally alter the conformational ensemble, but do alter interaction with the nucleic acid suggests an additional layer of complexity encoded in disordered proteins: on one side, the overall conformations of the protein may impact the capturing radius of the protein, whereas the specificity of residues in the sequence may modulate the binding affinity. This is particularly interesting since the properties of disordered regions can be robust to sequence mutations, as different residues can encode for similar properties of protein conformations, dynamics, and interactions. Indeed, available sequences of the SARS-CoV-2 genome are derived from patients and, therefore, are intrinsically biased to be functionally active (genome must be packaged and virus must be infective). Future studies will be required to understand what type of sequence mutations in IDRs can be tolerated by the virus to maintain the ability of condensing the nucleic acid.

### Conclusions

Overall, our measurements support a model in which the disordered NTD favors binding of the RNA to the RBD by directly participating in the interaction with the ligand and conformations are adapted based on the length of the nucleic acid. The dynamic nature of the complex combined with the preference of single-stranded RNAs may serve as a searching mechanism along the viral genome for identifying high affinity regions. The ability of the NTD domain to accommodate more than one RNA, possibly harnessing the hybridization of the sequence, may contribute to the packaging of the viral genome.

## Contributions

J.C. expressed, purified, and labeled all protein constructs. J.C. performed all single-molecule experiments with single-stranded RNA and folding stability of RBD_L_, including nanosecond FCS measurements. J.J.A. performed all single-molecule experiments with double-stranded RNA, folding stability measurements of RNA, and simulations. K.H. and J.J.A. *in-vitro* transcribed and purified RNA. J.C. and M.D.S.-B. designed the NTD-RBD_L_ and RBD_L_ nucleic acid binding assay. J.J.I. developed analytical tools for binding models. J.J.A. and A.S.H. developed computational tools for simulations. J.C., K.B.H, M.D.S.-B, J.J.I, J.J.A, A.S.H., and A.S. wrote the paper. M.D.S.-B. and A.S. supervised experiments and data analysis. J.J.I, J.C., M.D.S.-B., K.H., and A.S. conceived the experiments.

## Acknowledgements

We thank Tim Lohman and Roberto Galletto for useful insights and discussions on nucleic acid binding and SARS-CoV-2 Nucleocapsid, Ben Schuler and Daniel Nettels for developing and maintaining the Fretica package used for the analysis of single-molecule data, Silvia Jansen for sharing reagents, Vaclav Veverka and Evzen Boura for sharing their chemical shift perturbation data, and Giulio Tesei for useful insights regarding the simulation analysis. This research was supported by the NIH National Institute on Allergic and Infectious Diseases with R01AI163142 (to A.S., K.B.H., and A.S.H.) and by the NIH National Cancer Institute F99CA264413 (to J.J.A.). The content is solely the responsibility of the authors and does not necessarily represent the official views of the NIH.

