## Supplementary Information for "The disordered N-terminal tail of SARS CoV-2 Nucleocapsid protein forms a dynamic complex with RNA"

**Experimental setup and procedure for single-molecule fluorescence experiments.** Single-molecule confocal fluorescence measurements are performed on a Picoquant MT200 instrument (Picoquant, Germany). To enable Pulsed Interleaved Excitation (PIE), we synchronize a diode laser (LDH-D-C-485, PicoQuant, Germany) and a supercontinuum laser (SuperK Extreme, NKT Photonics, Denmark), filtered by a z582/15 band pass filter (Chroma) pulsed at 20 MHz such that a delay of approximately 25 ns occurs between each laser pulse. Lasers are focused in the sample through a 60x1.2 UPlanSApo Superapochromat water immersion objective (Olympus, Japan). Emitted photons are collected through the same objective, passed through a dichroic mirror (ZT568rpc, Chroma, USA), and further filtered by a long pass filter (HQ500LP, Chroma Technology) to suppress scattering light. After passing through the confocal pinhole (100 μm diameter), the emitted photons are separated into four channels by a polarizing beam splitter (which differentiates between perpendicular and parallel polarization), followed by a dichroic mirror (585DCXR, Chroma) that further discriminates between donor and acceptor photons. Donor and acceptor emission is then filtered using band pass filters, ET525/50m or HQ642/80m (Chroma Technology), respectively, and finally focused on SPAD detectors (Excelitas, USA). The arrival time of every photon is recorded with a HydraHarp 400 TCSPC module (PicoQuant, Germany). FRET experiments are performed by exciting the donor dye with a laser power of 100 μW (measured at the back aperture of the objective), whereas acceptor direct excitation is adjusted to match a total emission intensity after acceptor excitation to the one observed upon donor excitation (between 50 and 70 μW). Single-molecule FRET efficiency histograms are acquired at labeled protein concentrations between 50 pM and 100 pM, estimated from dilutions of samples with known concentration, as previously determined *via* absorbance measurements.

All measurements, unless differently specified, were performed in 50 mM Tris pH 7.4, 200 mM β-mercaptoethanol (for photoprotection), 0.001% Tween20 (for surface passivation) and

GdmCl at the reported concentrations. All measurements were performed in uncoated polymer coverslip cuvettes (Ibidi, Wisconsin, USA). When using denaturant or salt, the exact concentration is determined from measurement of the solution refractive index with an Abbe refractometer (Bausch & Lomb, USA).

Each sample was measured for at least 10 min at room temperature (295 ± 0.5 K) and all measurements were performed at least in duplicate (independent replicates from a new sample preparation) to confirm reproducibility of the results.

**Construction of transfer efficiency histograms.** Fluorescence bursts were identified by time-binning photons in bins of 1 ms and accepting bursts whose total number of photons after donor excitation was larger than at least 10 photons in each bin. Contiguous bins were merged if the total number of photons was larger than at least 20 photons. The exact threshold was selected based on the background contribution identified in the photon counting histograms with 1 ms binning. A minimum common threshold across constructs has been used to minimize variations in the width of the transfer efficiency distributions due to the difference in the acceptance thresholds, as expected for a shot-noise-limited system. Transfer efficiencies for each burst were calculated according to

$$E = n_A/(n_A + n_D) \qquad \textbf{Eq. S1}$$

where $n_A$ and $n_D$ are the numbers of donor and acceptor photons, respectively.

Reported transfer efficiencies are corrected for background, acceptor direct excitation, channel crosstalk, differences in detector efficiencies, and quantum yields of the dyes.

Similarly to transfer efficiency, the labeling stoichiometry ratio S is computed accordingly to:

$$S = I_D/(I_D + \gamma_{PIE} I_A) \qquad \textbf{Eq. S2}$$

where $I_D$ and $I_A$ represent the total intensities observed after donor and acceptor excitation and $\gamma_{PIE}$ provides a correction factor to account for the differences between donor and acceptor in detection efficiency and laser intensities. In the histograms, we present the bursts with stoichiometry corresponding to 1:1 donor:acceptor labeling (in contrast to donor and acceptor only populations), which are selected according to the criterion 0.3 < $S$ < 0.7. Variations in the selection criteria for the stoichiometry ratio do not impact significantly the observed mean transfer efficiency (within experimental errors).

**Fit of transfer efficiency distributions.** To estimate the mean transfer efficiency and extract multiple populations from the transfer efficiency histograms, each population was approximated with either a Gaussian or a LogNormal distribution function. When fitting more than one peak, the histogram is analyzed with a sum of Gaussian and/or LogNormal functions. When analyzing multiple overlapping populations, in order to limit the model parameters and potential overfitting, we favored the use of global fit analysis, where some parameters are shared across multiple or all concentrations.

**Folding equilibrium of the RBD.** The folding equilibrium of the RBD revealed the occurrence of three distinct states: native (N), intermediate (I), and unfolded (U). To quantify the thermodynamic properties of the three-state equilibrium N ⇌ I ⇌ U, the corresponding fraction folded, intermediate, and unfolded can be written in terms of the equilibrium constant $K_{UN}$ and $K_{NI}$ as:

$$f_U = 1/\left(1 + K_{UI} + K_{UI}K_{IN}\right) \quad \textbf{Eq. S3a}$$

$$f_I = K_{UI}/\left(1 + K_{UI} + K_{UI}K_{IN}\right) \quad \textbf{Eq. S3b}$$

$$f_N = K_{UI}K_{IN}/\left(1 + K_{UI} + K_{UI}K_{IN}\right) \quad \textbf{Eq. S3c}$$

The equilibrium constant $K_{UN}$ and $K_{NI}$ can be expressed as:

$$K_{UI} = exp\left[\Delta G_0^{UI}/RT\left(c - c^{UI}\right)/c^{UI}\right] \quad \textbf{Eq. S4a}$$

$$K_{IN} = exp\left[\Delta G_0^{IN}/RT\left(c - c^{IN}\right)/c^{IN}\right] \quad \textbf{Eq. S4b}$$

where $\Delta G_0^{UI}$ and $\Delta G_0^{IN}$ are the free energy differences in aqueous buffer conditions between the U and I and I and N states, respectively, and $c^{UI}$ and $c^{IN}$ are the concentrations where the corresponding fraction curves cross each other. It is important to note that whereas in the case of a simple two-state system N ⇌ U, the corresponding $c^{UN}$ represents the midpoint of the folding transition, in the general case with more than two states, the crossing points do not necessarily occur at the midpoint (50%) of the transition.

**Equilibrium binding models.** Here, we describe the assumptions behind the models for nonspecific interaction of monomeric N-protein with ssRNA. The models are derived for the specific case of the performed single-molecule experiments, where binding experiments were conducted at concentrations of protein much lower than the concentration of nucleic acid.

In all cases, we assume that binding takes place in a single orientation of the protein relative to the nucleic acid 3´-5´ polarity (though this can be easily extended to the more general case and does not significantly affect the interpretation of our results).

The observed association constants are expressed as the sum of intrinsic association constants for binding of the protein to each available position along the nucleic acid strand. We define a “position” as contiguous stretch of nucleotides that represent the protein’s binding footprint, i.e.:

$$K_A = \frac{\sum_i [(PR_M)_{position\ i}]}{[P][R_M]} = \sum_i K_{position\ i} \qquad \textbf{Eq. S5}$$

In these models, we assume that the oligonucleotides are homogeneous, made of repetitive superimposed segments presenting the same affinity for the protein, with periodicity length equal to 1 nucleotide; Coulombic end effects on counterion condensation and protein binding on the nucleic acid are not considered (see, for example, the work by Shkel, Ballin and Record [51])

The value of the intrinsic association constant $K_{int,m}$ for each available position is only dependent on the site size $m$ (i.e., the number of contiguous nucleotides involved in the interaction) but not on its position along the nucleic acid (we neglect end effects and position specificity). Under these assumptions the association constant can be written as

$$K_A = \frac{\sum_i [(PR_M)_{position\ i}]}{[P][R_M]} = \sum_i K_{position\ i} = \sum_{m=i}^{M} (\#\ positions\ with\ m\ cont.\ nt) K_{int,m} \qquad \textbf{Eq. S6}$$

*Single binding mode, no overhangs*

We first consider the case of a single binding mode with no overhangs. In this scenario:

- the protein only binds if the oligonucleotide length *M* is equal or longer than its contact site size, *n*; if *M* < *n*, it does not bind, i.e. $K_A = 0$;
- it binds with equal affinity, $K_{int}$, to all possible contiguous stretches of *n* nucleotides on the oligonucleotide, which can be counted to be *M* – *n* + 1;
- it does not bind through stretches of contiguous nucleotides shorter than the contact site size *n.*

Under these assumptions, the association constant can be written as:

$$K_A = \frac{\sum_i [(PR_M)_{position\ i}]}{[P][R_M]} = \sum_i K_{position\ i} = K_{int} 0 \quad for\ M < n \qquad \textbf{Eq. S7a}$$

$$K_A = \frac{\sum_i [(PR_M)_{position\ i}]}{[P][R_M]} = \sum_i K_{position\ i} = K_{int}(M - n + 1) \quad for\ M \geq n \qquad \textbf{Eq. S7b}$$

*Single binding mode, with 'overhangs'*

In this version of the model, the protein can bind to oligonucleotides of any length:

- if $M \geq n$, the protein can either bind in full length sites, spanning $n$ nucleotides, or to ends of the oligonucleotide, making contacts with a number of nucleotides smaller than $n$, leaving a protein 'overhang' that does not make contact with the nucleic acid;
- if $M < n$, the oligonucleotide can bind in different positions on the protein, spanning different portions of its nucleic acid binding site; these short oligos can bind within the binding site on the protein, or on the edges of the binding site leaving unbound nucleotide overhangs;
- in all cases, the protein interacts with a stretch of contiguous nucleotides, and the association constant for binding with a given number $m$ of contiguous nucleotides is equal to the product of an intrinsic association constant $K_{int,\ m}$, times the number of possible configurations for the given values of $M$ and $n$;
- the protein interacts with stretches of length $m < n$ only if there is no available nucleotides in one of the sides of the stretch; *i.e.*, only if binding to an end of an oligo or to an oligo with total length $M < n$;
- in addition to the fixed binding polarity, it is assumed that the nucleic acid binding site in the protein, able to interact with a contiguous stretch of nucleotides of length $n$, interacts with a short stretch of contiguous nucleotides, $m < n$, independently on where the stretch is located along the nucleic acid binding site; therefore,

$$K_A = K_{int,M}(n - M + 1) + 2 \sum_{j=1}^{M-1} K_{int,j} \quad for\, M < n \qquad \textbf{Eq. S8a}$$

$$K_A = K_{int}(M - n + 1) + 2 \sum_{j=1}^{n-1} K_{int,j} \quad for\, M \geq n \qquad \textbf{Eq. S8b}$$

- the values of intrinsic association constants with stretches of nucleotides shorter than *n*, $K_{int,\,m}$, are given by

$$K_{int,m} = K_{in,m} K_{tg} = (K_{in,n})^{\frac{m}{n}} K_{tg} = exp[\frac{\Delta G_{in,m} + \Delta G_{tg}}{RT}] = exp[\frac{(\Delta G_{in}/n)\, m + \Delta G_{tg}}{RT}] \qquad \textbf{Eq. S9}$$

The terms in the equation can be conceptualized with the following reaction scheme:

$$\mathrm{P} + \mathrm{R}_m \overset{K_{tg}}{\rightleftharpoons} \mathrm{P{\cdot}R}_m \overset{K_{in,m}}{\rightleftharpoons} \mathrm{PR}_m \qquad \textbf{Eq. S10}$$

where the first step is the bimolecular encounter of the protein P and the stretch of *m* nucleotides $R_m$, in the proper orientation for binding, and the second step is the actual establishment of the interactions between the protein and the nucleic acid site (compare with concepts derived by Lou and Sharp [52], equations 1-9)

This equation for $K_{int,m}$ involves the assumption that the translational-rotational entropic cost of the bimolecular encounter in the proper orientation ($\Delta G_{tg}$ = -RT Log$K_{tg}$) is independent of *m*. Also it involves the assumption that the contribution of the actual interactions established upon binding, -enthalpic and entropic components, such as counterion release- to the binding free energy is equally subdivided per nucleotide constituting the protein-nucleic acid binding interface ($\Delta G_{in,m} = m/n\ \Delta G_{in}$, or in terms of equilibrium constants, $K_{in,m} = (K_{in})^{m/n}$). Therefore:

$$K_A = K_{tg}\{(K_{in})^{M/n}(n - M + 1) + 2\sum_{j=1}^{M-1}(K_{in})^{j/n}\} \quad for\, M < n \qquad \textbf{Eq. S11a}$$

$$K_A = K_{tg}\{K_{in}(M - n + 1) + 2\sum_{j=1}^{n-1}(K_{in})^{j/n}\} \quad for\, M \geq n \qquad \textbf{Eq. S11b}$$

$$K_A = \frac{K_{int}}{K_{in}}\{(K_{in})^{M/n}(n - M + 1) + 2\sum_{j=1}^{M-1}(K_{in})^{j/n}\} \quad for\, M < n \qquad \textbf{Eq. S11c}$$

$$K_A = \frac{K_{int}}{K_{in}}\{K_{in}(M - n + 1) + 2\sum_{j=1}^{n-1}(K_{in})^{j/n}\} \quad for\, M \geq n \qquad \textbf{Eq. S11d}$$

where the first term represents the binding to the longest available stretch of nucleotides ($M$ if $M<n$, or $n$ if $M>n$) and the summation on the second term represents the binding to ends of the oligo with ends of the nucleic acid binding site on the protein involving shorter stretches of nucleotides.

The summation terms can be conveniently replaced by

$$\sum_{j=1}^{M-1}(K_{in})^{j/n} = (K_{in})^{M/n}\frac{1-(K_{in})^{(1-M)/n}}{(K_{in})^{1/n}-1} \qquad \textbf{Eq. S12a}$$

$$\sum_{j=1}^{n-1}(K_{in})^{j/n} = K_{in}\frac{1-(K_{in})^{(1-n)/n}}{(K_{in})^{1/n}-1} \qquad \textbf{Eq. S12b}$$

Then we have:

$$K_A = K_{int}\{(K_{in})^{M/n}[(n - M + 1) + 2\frac{1-(K_{in})^{(1-M)/n}}{(K_{in})^{1/n}-1}]\} \, for\, M < n \qquad \textbf{Eq. S13a}$$

$$K_A = K_{int}(M - n + 1) + 2\frac{1-(K_{in})^{(1-n)/n}}{(K_{in})^{1/n}-1} \, for\, M \geq n \qquad \textbf{Eq. S13b}$$

or, in terms of free energy,

$$K_A = K_{int} e^{\frac{\Delta G_{in}}{RT}(\frac{n-M}{n})} [(n - M + 1) + 2 \frac{e^{\frac{\Delta G_{in}}{RT}(\frac{M}{n})}(1-e^{\frac{\Delta G_{in}}{RT}(\frac{1-M}{n})})}{e^{\frac{\Delta G_{in}}{RT}(\frac{1}{n})}-1}] \, for \, M < n \quad \textbf{Eq. S14a}$$

$$K_A = K_{int} [(M - n + 1) + 2 \frac{e^{\frac{\Delta G_{in}}{RT}}(1-e^{\frac{\Delta G_{in}}{RT}(\frac{1-n}{n})})}{e^{\frac{\Delta G_{in}}{RT}(\frac{1}{n})}-1}] \, for \, M \geq n \quad \textbf{Eq. S14b}$$

**Nanosecond FCS analysis.** Autocorrelation curves of acceptor and donor channels and cross-correlation curves between acceptor and donor channels were calculated with the methods described previously [53,54]. All samples were measured at single-molecule concentrations (~100 pM), and bursts corresponding to the donor-acceptor population transfer efficiency were selected to eliminate the contribution of donor-only to the correlation amplitude. Finally, the correlation was computed over a time window of 5 μs, and characteristics timescales were extracted according to:

$$g_{ij}(\tau) = 1 + \frac{1}{N}(1 - c_{AB} Exp[-(\tau - \tau_0)/\tau_{AB}]) \times$$

$$\times (1 + c_{CD} Exp[-(\tau - \tau_0)/\tau_{CD}])(1 + c_T Exp[-(\tau - \tau_0)/\tau_T]) \quad \textbf{Eq S15}$$

where *N* is the mean number of molecules in the confocal volume and $i$ and $j$ indicate the type of signal (either from the Acceptor or Donor channels). The three multiplicative terms describe the contribution to amplitude and timescale of photon antibunching (AB), chain dynamics (CD), and triplet blinking of the dyes (T). $\tau_{CD}$ is then converted in the reconfiguration time of the interdye distance $\tau_r$ correcting for the filtering effect of FRET as described previously [55].

**Coarse-grained simulations**

Coarse-grained simulations were performed using the Mpipi model[28]. In Mpipi, each bead (amino acid or nucleotide) is chemically unique, and inter-bead interactions contain contributions from a short-range Wang-Frenkel potential and, where applicable, a long-range Coulombic potential for beads with a net charge[56]. The Coulombic potential takes the ionic strength into account, and simulations were performed at an equivalent of 50 mM NaCl. The parameters associated with the inter-bead Wang-Frenkel potential were determined through a combination of all-atom and quantum mechanical simulations and capture a mixture of Van der Waal interactions, cation-pi and pi-pi interactions.

As in previous work, folded domains were modeled as rigid bodies, whereas intrinsically disordered regions and ssRNA were described as flexible polymers[28,29]. Beads found within the core of globular domains ("buried" residues) have their interaction strength scaled down, as in the original Mpipi implementation.

*Calculating apparent association constants from simulations*

To determine the apparent association constants ($K_A$) for simulations, we calculated the fraction of frames in which protein and RNA were bound. To determine the bound fraction requires a definition for protein:RNA binding. We applied a measure whereby binding is determined based on consecutive simulation frames in which the protein and RNA centers-of-mass (COM) are under an RNA-length dependent threshold. This approach is motivated by the fact that histograms of the protein:RNA COM clearly show two distributions; a bound COM distance distribution and an unbound COM distance distribution (**Supplementary Fig. 5B**). As the RNA becomes longer, the separation between these two peaks changes (as the peak of the bound distribution shifts to larger values due to the larger RNA molecule). These histograms enable us to define an RNA-length-specific distance threshold for each simulation. With this naive cutoff defined, we define binding as five or more consecutive frames where the protein and RNA COM are under the predefined threshold distance. The use of a minimum number of consecutive frames enables us to distinguish transient random encounters between the protein and RNA from *bona fide* binding events, where protein and RNA are directly engaging (**Supplementary Fig. 5C,D)**.

Having determined the fraction bound, we then calculated an apparent $K_D$ with the expression:

$$K_D = \frac{(1-f_{bound})^2}{N_A V f_{bound}} \quad \textbf{Eq. S16}$$

where $f_{bound}$ is the fraction of the simulation in which the two species are bound, $N_A$ is Avogadro's constant, and V is the simulation box volume in liters, returning a $K_D$ in mol/L. The $K_A$ is then calculated as 1 / $K_D$. This approach is analogous to that of Tesei *et al.*, albeit using a different strategy to define if two molecules are bound vs. unbound[57]. Finally, having calculated the apparent association constants, we can ask how protein:RNA affinity varies across simulations of the NTD alone, RBD alone, and NTD-RBD with different lengths of $(rU)_n$.

When comparing the $K_A$ values from simulations with experiment, we found poor agreement between the absolute values of the association constants, a feature that is commonly seen for coarse-grained models[58]. To enable a direct comparison between experiments and simulations, we calculate a normalized binding affinity ($K_A^*$), which we define as the ratio between the simulation (or experimental) apparent $K_A$ for a given protein:RNA combination divided by the corresponding simulation (or experimental) apparent $K_A$ for NTD-RBD binding to $(rU)_{25}$. This, in effect, sets the NTD-RBD + $(rU)_{25}$ binding affinity as a reference point, and all other $K_A^*$ values are defined as either greater than 1 (stronger binding than NTD-RBD + $(rU)_{25}$) or less than 1 (weaker binding than NTD-RBD + $(rU)_{25}$). By using this ratio, we can plot data from simulations and experiments on the same axes and compare the relative binding affinities (as a function of RNA length, protein construct, or protein sequence). This analysis reveals relatively good agreement between simulations and experiments (**Fig. 5D, F, G**), despite the many assumptions made in the coarse-grained force field.

#### Measuring the stability of double-stranded RNA

Absorbance is measured using a UV-Vis spectrophotometer. The sharp increase in absorbance reports on the hyperchromicity of the hairpin RNA as it converts from double-stranded (ds) to single-stranded (ss) RNA. Melting temperatures are determined by fitting absorbance values as a function of temperature to:

$$Abs = \frac{(\alpha_{ds} + \beta_{ds}T) + (\alpha_{ss} + \beta_{ss}T)e^{-m(T-T_m)}}{1 + e^{-m(T-T_m)}} \quad \textbf{Eq. S17}$$

where $\alpha_{ds}$ and $\alpha_{ss}$ refer to the absorbance of the RNA in the ds and ss state at initial temperature. $\beta_{ds}$ and $\beta_{ss}$ are the rate of change of the absorbance in each state as a function of temperature (*T*) in Kelvin. $m$ is the $m$-value and $T_m$ is the temperature at the midpoint of the transition from ds to ss RNA.

### SUPPLEMENTARY FIGURES.

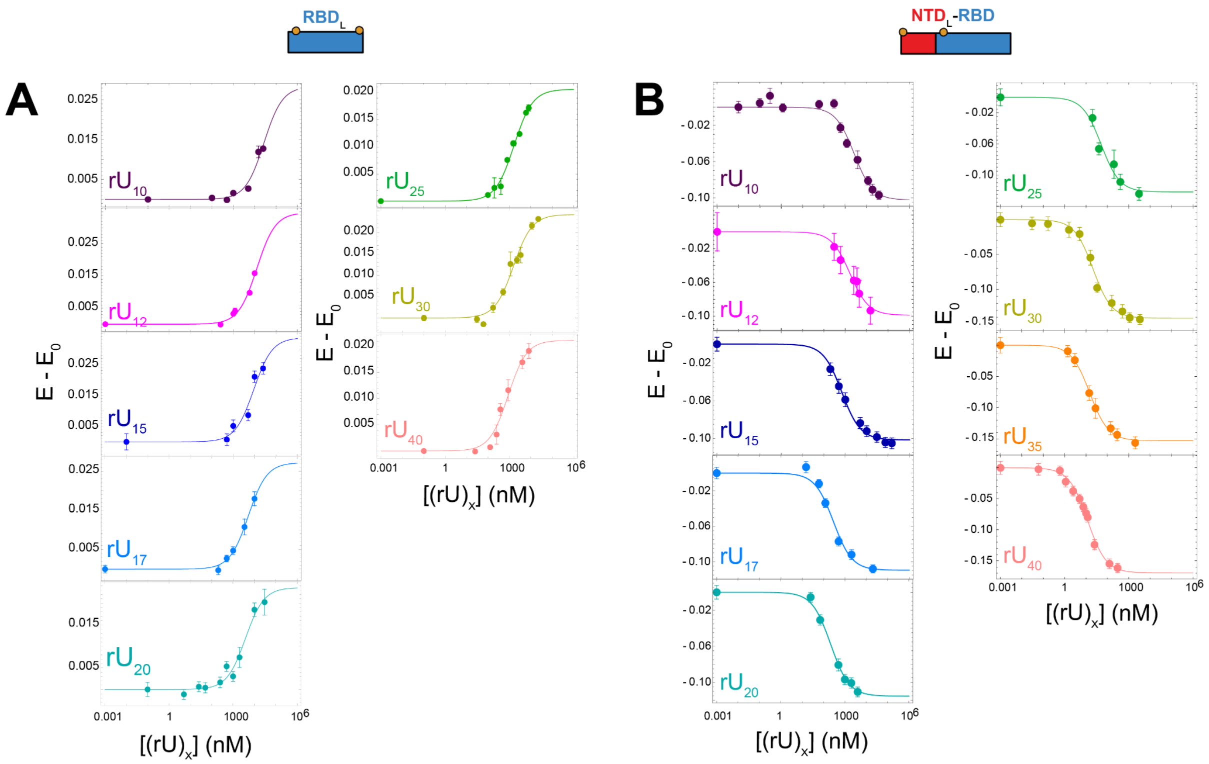


**Supplementary Figure 1.** Transfer efficiency variation upon binding for RBD$_L$ (panel **A**) and NTD$_L$-RBD (panel **B**). Error bars represent the standard deviation of at least two independent experiments. Solid lines are fit to **Eq. 1.**

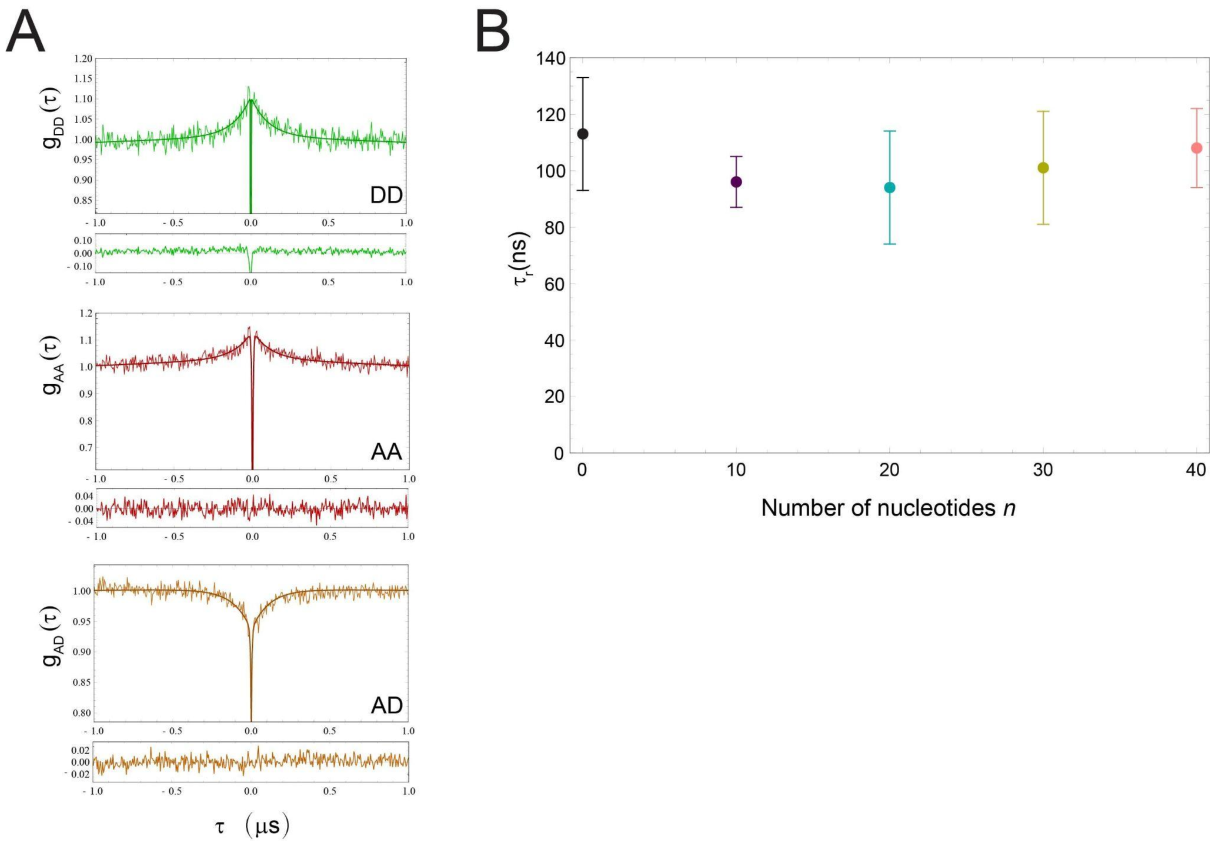


**Supplementary Figure 2. Dynamics of the disordered NTD when complex with RNA. A.** Example of nanosecond-FCS (nsFCS) traces of NTD$_L$-RBD in the presence of (rU)$_{40}$. Donor-donor, acceptor-acceptor, and donor-acceptor correlations are shown in green, red, and orange (respectively) with the fit according to **Eq. S15** and corresponding residuals. **B.** Reconfiguration times computed for the chain in the absence and in the presence of (rU)$_n$ with *n* = 10, 20, 30, 40.

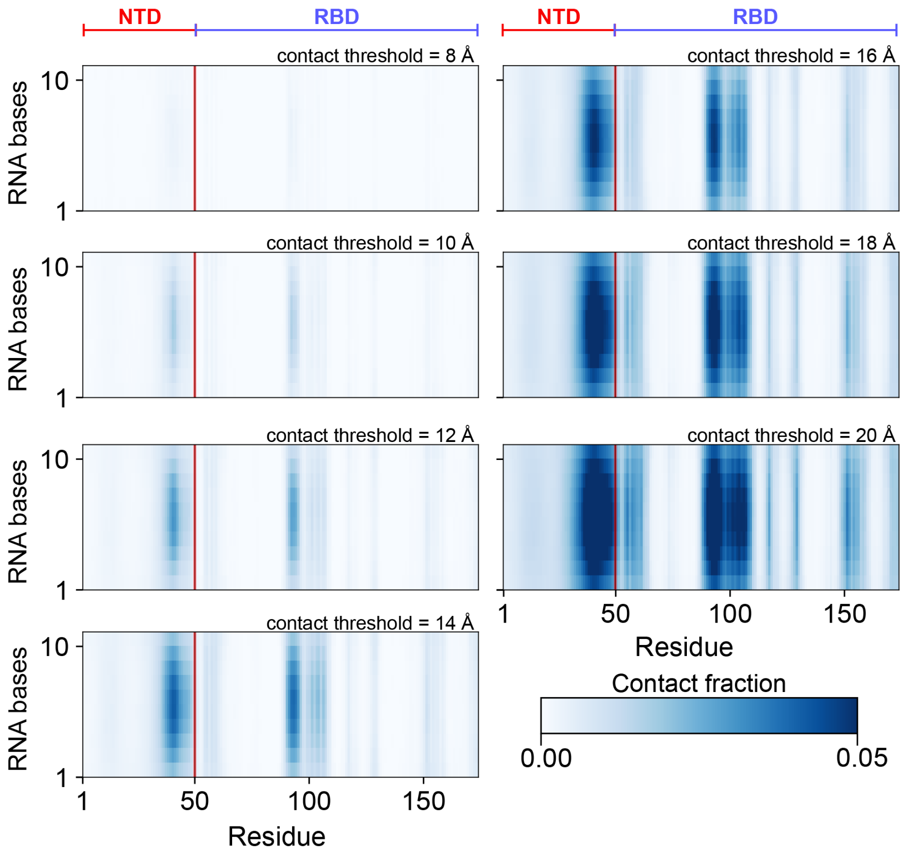


**Supplementary Figure 3. NTD-RBD:(rU)$_{10}$ dependence of interacting residues on distance threshold used for contact fraction.** The distance threshold used to define nucleotide:amino acid contacts was varied from 8 Å to 20 Å to assess how this altered the residues identified as RNA interacting. While, as expected, the contact fraction systematically changes as the threshold increases, the pattern of residues engaging in protein:RNA interactions remains consistent.

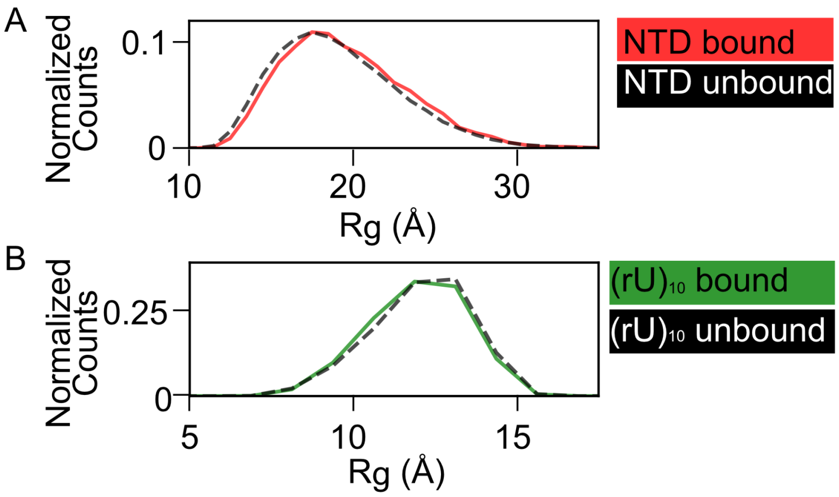


**Supplementary Figure 4. The NTD and RNA remain disordered in the NTD-RBD:$(rU)_{10}$ complex**. **A.** Histogram showing the radius of gyration ($R_g$) distribution for the NTD region taken from either the NTD-RBD:$(rU)_{10}$ complex (red line) or from the unbound (black line) states of NTD-RBD. The relative histogram counts have been normalized to the total number of events in the bound or unbound state. Specifically, 8,198 frames were RNA-bound in the trajectory analyzed, while 81,347 were RNA-unbound. If the NTD folded upon binding, we would expect to see a tighter distribution for the $R_g$ at a smaller mean value, yet the $R_g$ distribution in the bound state remains broad, with a slightly smaller mean value in the unbound state reflective of the length dependent expansion of the NTD upon binding (unbound NTD$\langle R_g \rangle$= 19.1 Å, bound NTD$\langle R_g \rangle$= 19.6 Å). The root-mean-square value of the end-to-end distance is reported in **Fig. 5C**. **B.** Analogous analysis from the perspective of the $(rU)_{10}$. The mean value is similar in the bound vs. unbound states (unbound $(rU)_{10}\langle R_g \rangle$= 10.6 Å, bound $(rU)_{10}\langle R_g \rangle$= 10.7 Å), but the broad distribution remains consistent with a largely disordered ensemble of conformations. See also **Supplementary Movie S1.**

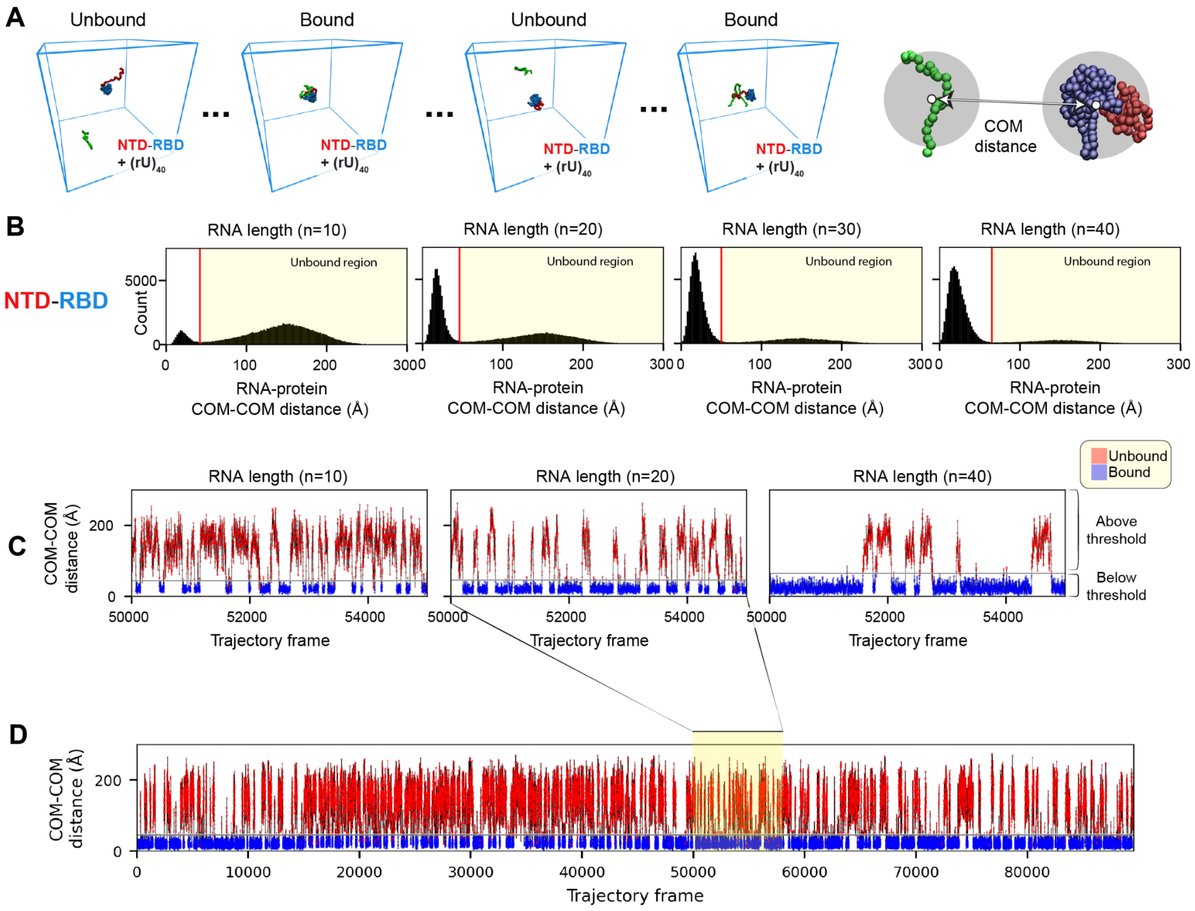


**Supplementary Figure 5. Simulations of N protein construct and RNA binding. A.** Example simulation snapshots from the NTD-RBD + $(rU)_{40}$ simulation showing bound and unbound configurations. On the far right a schematic of the center of mass (COM) distance is shown for NTD-RBD and $(rU)_{25}$ that are 101 Å apart. The COM for each of the two molecules is calculated using the get_center_of_mass() function in SOURSOP. **B.** Intermolecular center-of-mass (COM) distance between the protein and RNA molecules enables us to define a distance threshold that can be used to define when the two molecules are bound vs. unbound. The distance threshold for NTD-RBD binding to RNA varies between 42 Å (for RNA of length 10) and 65 Å (for RNA of length 40). Note that this distance reflects the center of mass between the two molecules, not the minimum distance. **C.** Subtrajectory taken from a simulation showing bound and unbound states being automatically delineated based on the combination of the distance threshold introduced in panel A, alongside the requirement for five or more consecutive frames under the cutoff threshold to be used to define binding (or lack thereof). Panel C shows sub-trajectories from simulations with RNAs of length 10, 20, and 40 nucleotides. **D.** Full trajectory of simulations with RNA of length 20 showing over 200 independent binding and unbinding events for each RNA length.

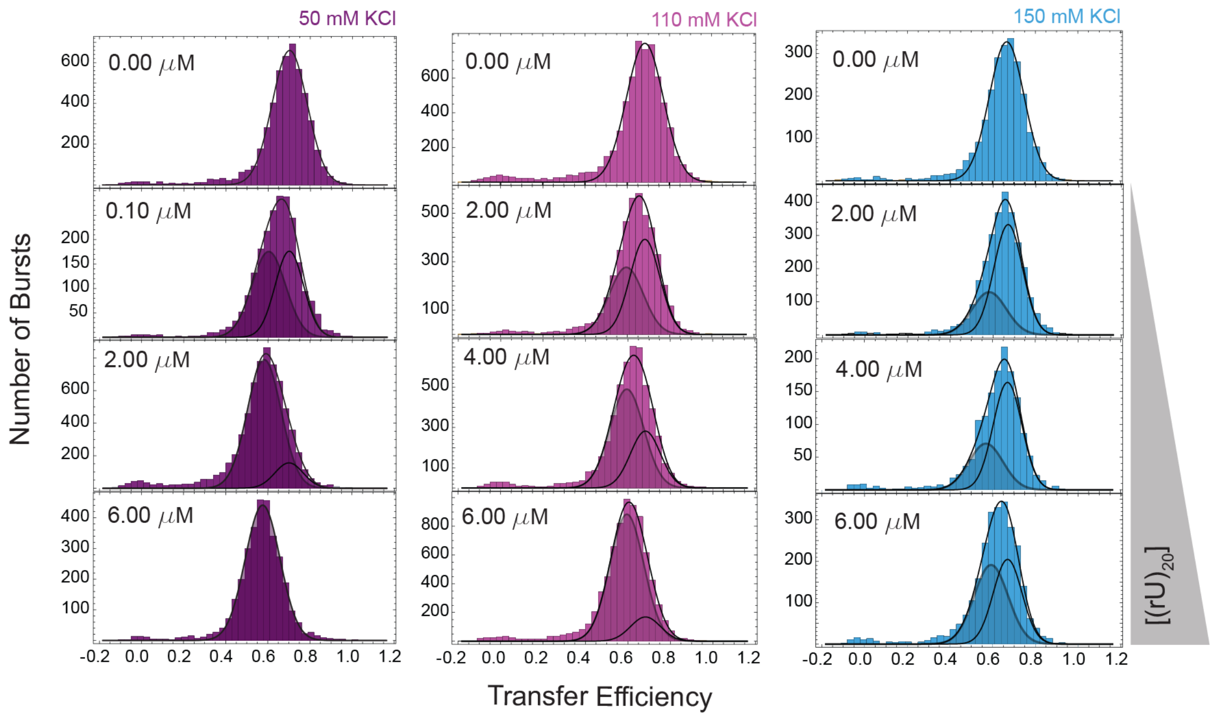


**Supplementary Figure 6. Representative transfer efficiency distributions of $(rU)_{20}$ as a function of salt concentration.** Histograms of transfer efficiencies measured at 50 mM KCl (left, purple), 110 mM KCl (center, magenta), 150 mM KCl (right, blue) from 0 to 6 μM $(rU)_{20}$.

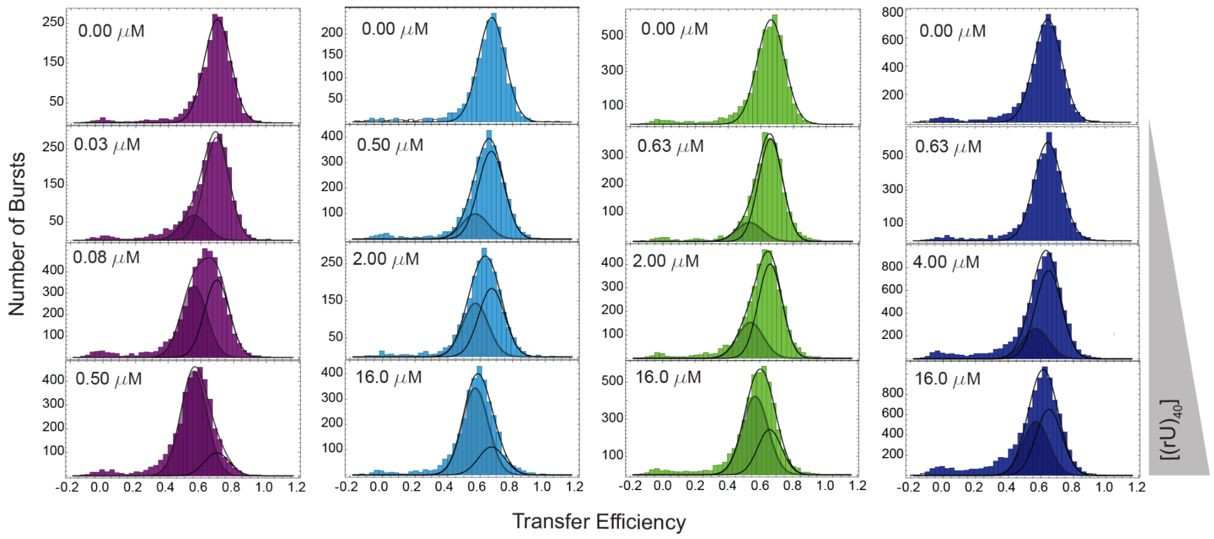


**Supplementary Figure 7. Representative transfer efficiency distributions of $(rU)_{40}$ as a function of salt concentration.** Histograms of transfer efficiencies measured at 50 mM KCl (left, purple), 110 mM KCl (center, magenta), 150 mM KCl (right, blue) from 0 to 16 μM $(rU)_{40}$. Distributions are fitted with up to two Gaussian distributions to quantify the fraction bound and unbound and the corresponding transfer efficiencies.

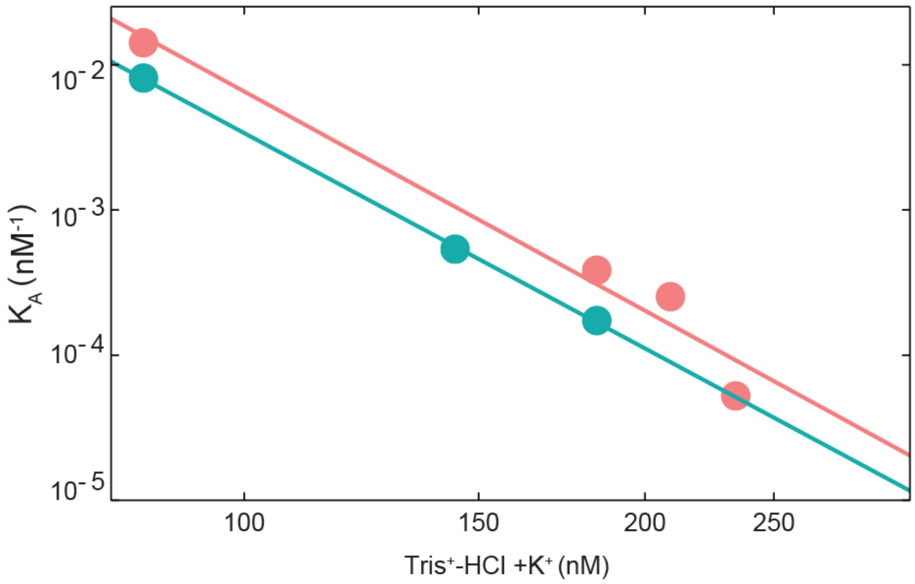


**Supplementary Figure 8. Association constant as a function of the total concentration of positive ions for $(rU)_{20}$ (cyan) and $(rU)_{40}$ (pink).** Errors associated with each $K_A$ are standard errors of the fit and are reported in **Supplementary Table 12** (not visible because smaller than the marker for the experimental point).

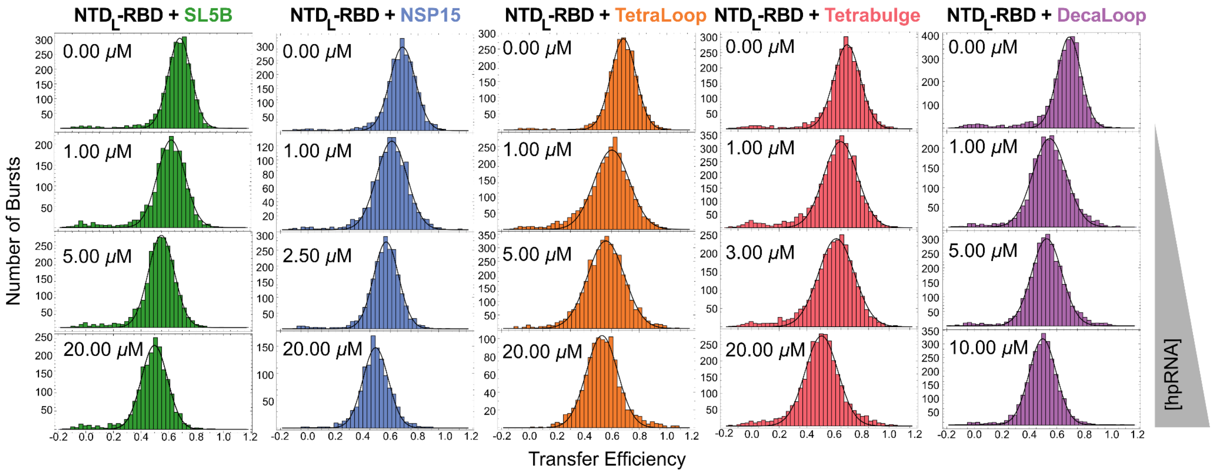


**Supplementary Figure 9. Transfer efficiency distributions for NTD$_L$-RBD and RNA hairpins.** Representative histograms of NTD$_L$-RBD + hairpin RNA (hpRNA) as a function of concentration .

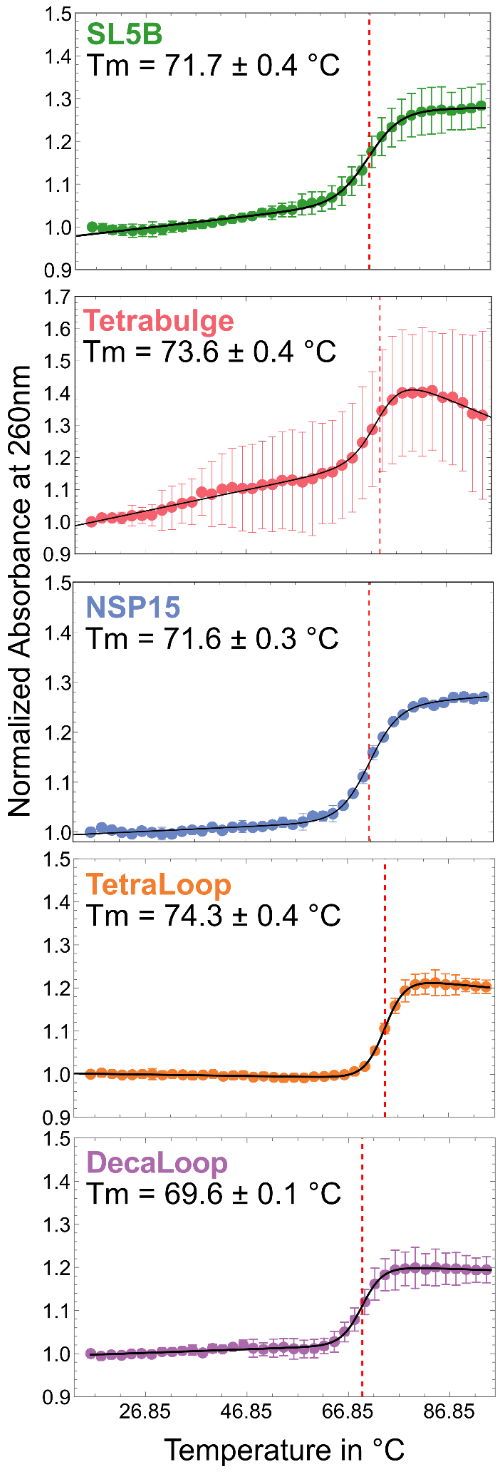


**Supplementary Figure 10. Thermal melting curves of RNA hairpins.** Absorbance at 260 nm was monitored over a temperature range of 16 °C to 95 °C in 10 mM HEPES, 50 mM KCl, 0.5 mM EDTA, pH 7.4 (23 °C). Temperature was increased in 2 °C steps at a rate of 1 °C/minute and data collected for 2 s after equilibration for 2 minutes after each step. Dots and error bars represent the mean and standard error of 2 measurements performed on different samples. Black solid lines are simulations of Eq. S17 fitted to the data by least squares nonlinear regression. The best fit value plus/minus the standard error of the fit for the $T_m$ is shown in the plots.

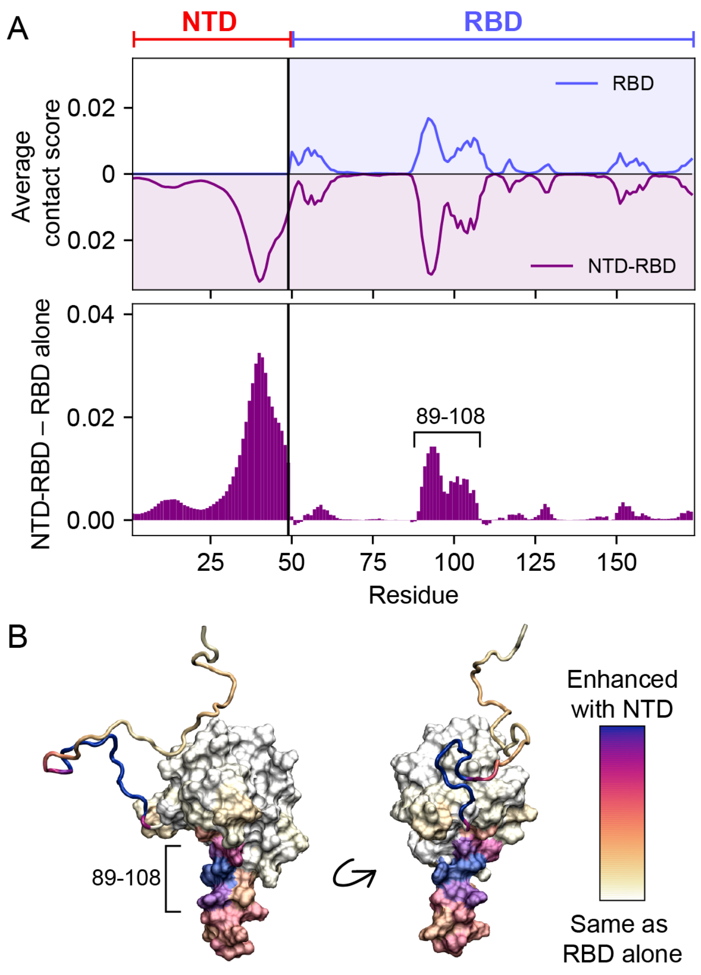


**Supplementary Figure 11. The NTD does not alter the overall pattern of RBD:RNA interactions. A.** To easily compare RBD:RNA interactions with and without the NTD, we calculated the average per-residue contact score for NTD-RBD + $(rU)_{10}$ and RBD + $(rU)_{10}$. Specifically, this involved averaging the per-residue contact fraction over the ten nucleotides to give a per-nucleotide interaction score (which we define as the average contact score). The scores for RBD alone vs. NTD-RBD are shown above. The profiles effectively mirror one another, even down to fine detail, supporting the notion that in our simulations, the addition of the NTD does not alter which residues on the RBD RNA interact with. However, the frequency with which specific sub-regions interact with RNA does change upon the addition of the NTD. Notably, by comparing the difference in average scores (i.e., NTD-RBD – RBD, bottom panel), residues 89 – 108 within the RBD show an uptick in RNA contacts. **B.** We annotated a structural model of the NTD-RBD by coloring residues according to their enhanced RNA interaction in the presence of the NTD (i.e., scores shown in the bottom panel of panel A). This annotation clearly shows residues in the β-extension dominate in terms of the NTD-enhanced RNA binding.

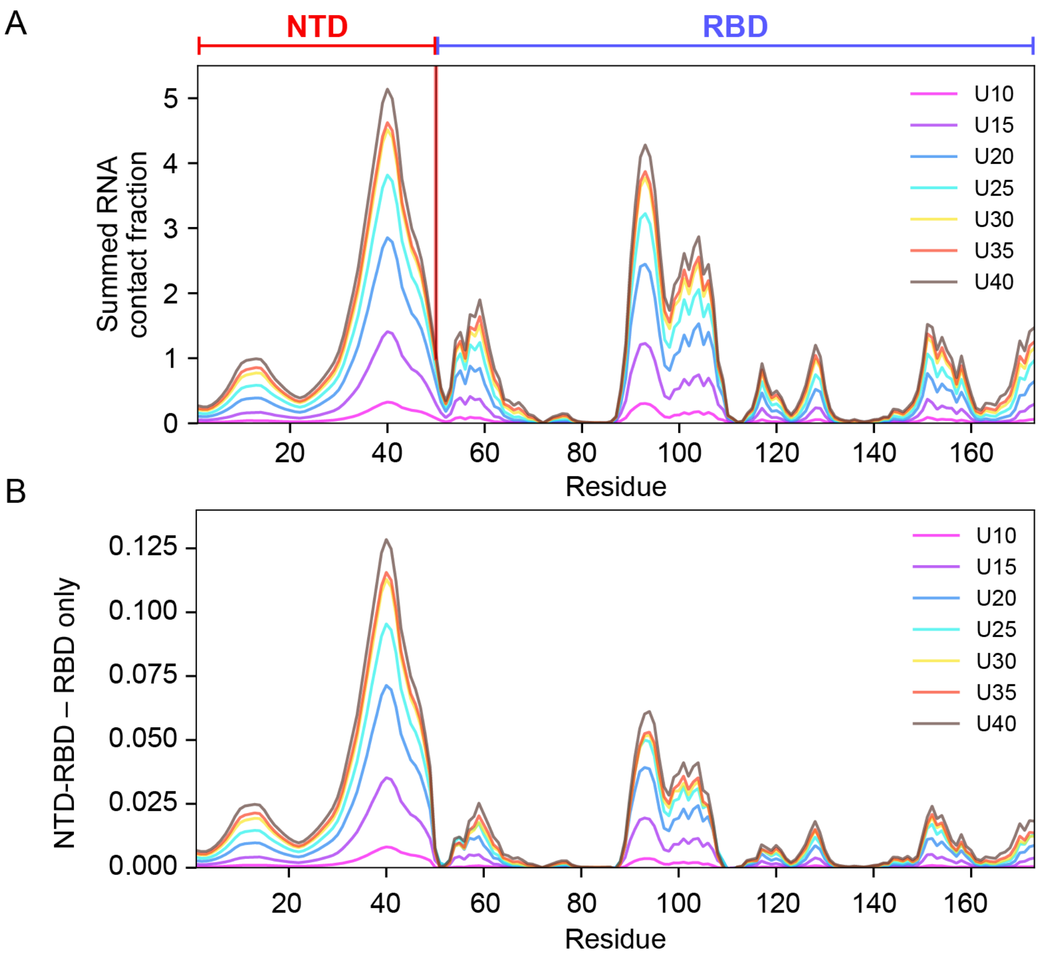


**Supplementary Figure 12. RNA length tunes the magnitude of protein:RNA interactions but does not alter the overall pattern of RBD:RNA interactions**. **A.** Following the analysis in **Supplementary Fig. 11A**, we calculated the summed contact fraction for each residue across $(rU)_{10}$, $(rU)_{15}$, $(rU)_{20}$, $(rU)_{25}$, $(rU)_{30}$, $(rU)_{35}$, and $(rU)_{40}$. By comparing these profiles, our analysis reveals that as the rU becomes longer, the regions identified in our initial analysis (residue 30–50 and residues 89 – 109) show an RNA-length-dependent enhancement in protein:RNA contacts, supporting the interpretation that these two regions are the primary determinants of protein:RNA interaction. Outside of these regions, additional loci on both the NTD and RBD also engage with RNA in an RNA-length-dependent manner. In all cases, contacts observed in NTD-RBD:$(rU)_{10}$

simulations (**Supplementary Fig. 11A**) were enhanced as a function of RNA length, but we did not observe novel interactions appear with longer RNA molecules. **B.** We assessed how the presence of the NTD altered RBD:RNA interaction by subtracting RBD contact fractions from NTD-RBD contact fractions across the same six RNA lengths. This analysis confirmed conclusions drawn for using $(rU)_{10}$ – that the major subregion within the RBD that is influenced by the presence of the NTD is the positively-charged β-extension (specifically in residues 89–108) (**Supplementary Fig. 11B**).

**Supplementary Movie 1: NTD and RNA remain disordered in the bound state.**

Movie of NTD-RBD + $(rU)_{25}$ showing frames from a bound-state ensemble.

**Supplementary Table 1. Sequence of wild type NTD-RBD.** Labeling positions are reported in red.

| | |
|---|---|
| 1 | MSDNGPQNQR NAPRITFGGP SDSTGSNQNG ERSGARSKQR RPQGLPNNTA |
| 51 | SWFTALTQHG KEDLKFPRGQ GVPINTNSSP DDQIGYYRRA TRRIRGGDGK |
| 101 | MKDLSPRWYF YYLGTGPEAG LPYGANKDGI IWVATEGALN TPKDHIGTRN |
| 151 | PANNAAIVLQ LPQGTTLPKG FYA |

**Supplementary Table 2. Constructs used in this study.** For each construct, we reported the start and end positions compared to the wild type (WT) sequence, the labeling positions, and highlighted in yellow the portion of the sequence in between the labeling positions.

| Name | Sequence | Start Position (WT) | End Position (WT) | Labeling Positions (WT) |
|---|---|---|---|---|
| **$NTD_L$-RBD** | GPCSDNGPQNQRNAPRITFGGPSDSTGSNQNGERSGARSKQRRPQGLPNNTASWFTALTQHGKEDLKFPCGQGVPINTNSSPDDQIGYYRRATRRIRGGDGKMKDLSPRWYFYYLGTGPEAGLPYGANKDGIIWVATEGALNTPKDHIGTRNPANNAAIVLQLPQGTTLPKGFYA | **1** | **173** | **1, 68** |

| | | | | |
|---|---|---|---|---|
| **NTD$_L$-RBD**<br>**Omicron**<br>**(P13L, Δ31-33)** | GPCSDNGPQNQRNALRITFGGPSDS<br>TGSNQNGGARSKQRRPQGLPNNTAS<br>WFTALTQHGKEDLKFPCGQGVPINT<br>NSSPDDQIGYYRRATRRIRGGDGKM<br>KDLSPRWYFYYLGTGPEAGLPYGAN<br>KDGIIWVATEGALNTPKDHIGTRNP<br>ANNAAIVLQLPQGTTLPKGFYA | **1** | **173** | **1, 68** |
| **NTD-RBD$_L$** | GPMSDNGPQNQRNAPRITFGGPSDS<br>TGSNQNGERSGARSKQRRPQGLPNN<br>TASWFTALTQHGKEDLKFPCGQGVP<br>INTNSSPDDQIGYYRRATRRIRGGD<br>GKMKDLSPRWYFYYLGTGPEAGLPY<br>GANKDGIIWVATEGALNTPKDHIGT<br>RNPANNAAIVLQLPQGTTLPKGFCA | **1** | **173** | **68,172** |
| **RBD$_L$** | GPGLPNNTASWFTALTQHGKEDLKF<br>PCGQGVPINTNSSPDDQIGYYRRAT<br>RRIRGGDGKMKDLSPRWYFYYLGTG<br>PEAGLPYGANKDGIIWVATEGALNT<br>PKDHIGTRNPANNAAIVLQLPQGTT<br>LPKGFCA | **44** | **173** | **68,172** |

**Supplementary Table 3. RNA sequences used in this study.**

| RNA | Genomic position | Nts | Sequence 5’-3’ | Origin |
|---|---|---|---|---|
| **Poly(rU)** | - | <250 | | Midland Certified Reagent Company |
| **Poly(rU)** | - | 10,12,15, 17,20,25, 30,35,40 | | IDT, Horizon Discovery |
| **V21** | 127-148 | 21 | **UAUAAUUAAUAACUA AUUACU** | IDT, Horizon Discovery |
| **SL5B*** | 228 - 252 | 30 | **GGGCAUACCUAGGU UUCGUCCGGGUGUG CC** | *in vitro transcribed* |
| **NSP15** | 19972-20000 | 31 | **GGGCUCACUGUCUU UUUUGAUGGUAGAG UCC** | *in vitro transcribed* |
| **Tetraloop** | based on NSP15 | 29 | **GGGCUCACUGUCUU CGGAUGGUGAGCUC** | *in vitro transcribed* |
| **Decaloop** | based on NSP15 | 34 | **GGGCUCACUGUCUU CUUUUUUUGAUGGU GAGCUC** | *in vitro transcribed* |
| **Tetrabulge** | based on NSP15 | 30 | **GGGCUCACUGUCUU CGGAUGGUAGAGUC C** | *in vitro transcribed* |

*The SL5B sequence is taken from the SARS-CoV genome and differs for two nucleotides from the SARS-CoV-2 genome.

**Supplementary Table 4. Summary of simulation details.**

| Simulation components | Box size ($nm^3$) | Temp (K) | Steps per simulation (millions) | Number of independent replicas | Total production frames |
|---|---|---|---|---|---|
| NTD | 30 | 298 | 300 | 30 (5x6 starting conf) | 268,650 |
| RBD | 30 | 298 | 300 | 30 (5x6 starting conf) | 268,420 |
| NTD-RBD | 30 | 298 | 300 | 25 (5x5 starting conf) | 221,592 |
| NTD + $(rU)_{25}$ | 30 | 298 | 300 | 30 (5x6 starting conf) | 268,650 |
| RBD + $(rU)_{10}$ | 30 | 298 | 300 | 30 (5x6 starting conf) | 268,650 |
| RBD + $(rU)_{12}$ | 30 | 298 | 300 | 30 (5x6 starting conf) | 268,650 |
| RBD + $(rU)_{15}$ | 30 | 298 | 300 | 30 (5x6 starting conf) | 264,405 |
| RBD + $(rU)_{17}$ | 30 | 298 | 300 | 30 (5x6 starting conf) | 268,322 |
| RBD + $(rU)_{20}$ | 30 | 298 | 300 | 30 (5x6 starting conf) | 267,693 |
| RBD + $(rU)_{25}$ | 30 | 298 | 300 | 30 (5x6 starting conf) | 265,683 |
| RBD + $(rU)_{30}$ | 30 | 298 | 300 | 30 (5x6 starting conf) | 265,026 |
| RBD + $(rU)_{35}$ | 30 | 298 | 300 | 30 (5x6 starting conf) | 268,650 |
| RBD + $(rU)_{40}$ | 30 | 298 | 300 | 30 (5x6 starting conf) | 263,567 |
| RBD + $(rU)_{180}$ | 30 | 298 | 300 | 30 (5x6 starting conf) | 268,503 |
| NTD-RBD + $(rU)_{10}$ | 30 | 298 | 300 | 30 (5x6 starting conf) | 268,650 |
| NTD-RBD + $(rU)_{12}$ | 30 | 298 | 300 | 30 (5x6 starting conf) | 268,650 |
| NTD-RBD + $(rU)_{15}$ | 30 | 298 | 300 | 30 (5x6 starting conf) | 267,563 |
| NTD-RBD + $(rU)_{17}$ | 30 | 298 | 300 | 30 (5x6 starting conf) | 267,341 |

| | | | | | |
|---|---|---|---|---|---|
| NTD-RBD + $(rU)_{20}$ | 30 | 298 | 300 | 30 (5x6 starting conf) | 265,965 |
| NTD-RBD + $(rU)_{25}$ | 30 | 298 | 300 | 30 (5x6 starting conf) | 263,434 |
| NTD-RBD + $(rU)_{30}$ | 30 | 298 | 300 | 30 (5x6 starting conf) | 268,134 |
| NTD-RBD + $(rU)_{35}$ | 30 | 298 | 300 | 30 (5x6 starting conf) | 268,650 |
| NTD-RBD + $(rU)_{40}$ | 30 | 298 | 300 | 30 (5x6 starting conf) | 267,220 |
| NTD-RBD + $(rU)_{180}$ | 30 | 298 | 300 | 30 (5x6 starting conf) | 268,650 |
| OmNTD-RBD (P13L,$\Delta^{31\text{-}33}$) + $(rU)_{25}$ | 30 | 298 | 300 | 30 (5x6 starting conf) | 267,333 |
| OmNTD-RBD (P13L) + $(rU)_{25}$ | 30 | 298 | 300 | 30 (5x6 starting conf) | 268,527 |
| OmNTD-RBD ($\Delta^{31\text{-}33}$) + $(rU)_{25}$ | 30 | 298 | 300 | 30 (5x6 starting conf) | 266,023 |

**Supplementary Table 5. RBD Folding parameters.**

| | $RBD_L$ full-length protein | $RBD_L$ |
|---|---|---|
| $c_{1/2,IF}$ (M) | 1.68 ± 0.02 | 1.26 ± 0.03 |
| $c_{1/2,IF}$ (M) | 1.64 ± 0.02 | 1.21 ± 0.01 |
| $\Delta G_{UI}$ (kcal $mol^{-1}M^{-1}$) | 6.6 ± 0.5 | 2.8 ± 0.1 |
| $\Delta G_{IF}$ (kcal $mol^{-1}M^{-1}$) | 8.1 ± 0.5 | 7.6 ± 0.4 |

**Supplementary Table 6. Intrinsic association constants**

| | $K_{int}$ for poly(rU) ($\mu M^{-1}$) nucleotides |
|---|---|
| $RBD_L$ | (6 ± 1) x $10^{-2}$ |
| NTD-$RBD_L$ | 2.0 ± 0.4 |
| $NTD_L$-RBD | 4.0 ± 0.3 |

**Supplementary Table 7. $RBD_L$ and $NTD_L$-RBD association constants for $(rU)_n$** as measured by single-molecule FRET

| | $K_A$ ($\mu M^{-1}$) molecules | |
|---|---|---|
| n | $RBD_L$ | $NTD_L$-RBD |
| 10 | $(4 \pm 3) \times 10^{-2}$ | $(3.8 \pm 0.1) \times 10^{-1}$ |
| 12 | $(7.9 \pm 0.4) \times 10^{-2}$ | $(6.1 \pm 1.5) \times 10^{-1}$ |
| 15 | $(1.1 \pm 0.6) \times 10^{-1}$ | $1.4 \pm 0.1$ |
| 17 | $(1.9 \pm 0.4) \times 10^{-1}$ | $3.5 \pm 0.8$ |
| 20 | $(2.6 \pm 0.7) \times 10^{-1}$ | $4.6 \pm 0.6$ |
| 25 | $(6.0 \pm 0.9) \times 10^{-1}$ | $20 \pm 5$ |
| 30 | $(6.6 \pm 1.3) \times 10^{-1}$ | $47 \pm 6$ |
| 35 | - | $66 \pm 5$ |
| 40 | $1.2 \pm 0.3$ | $85 \pm 7$ |

**Supplementary Table 8. Simulation-derived association constants ($K_A$) in $\mu M^{-1}$**

| **Construct** | **NTD;** $K_A$ ($\mu M^{-1}$) | **RBD;** $K_A$ ($\mu M^{-1}$) | **NTD-RBD;** $K_A$ ($\mu M^{-1}$) |
|---|---|---|---|
| $(rU)_{10}$ | $(9.5 \pm 0.7) \times 10^{-5}$ | $(4.6 \pm 0.4) \times 10^{-4}$ | $(1.90 \pm 0.05) \times 10^{-3}$ |
| $(rU)_{12}$ | $(2.5 \pm 0.2) \times 10^{-4}$ | $(8.7 \pm 0.8) \times 10^{-4}$ | $(4.2 \pm 0.1) \times 10^{-3}$ |
| $(rU)_{15}$ | $(5.6 \pm 0.4) \times 10^{-4}$ | $(1.7 \pm 0.1) \times 10^{-3}$ | $(1.00 \pm 0.03) \times 10^{-2}$ |
| $(rU)_{17}$ | $(8.0 \pm 0.5) \times 10^{-4}$ | $(2.5 \pm 0.2) \times 10^{-3}$ | $(1.8 \pm 0.1) \times 10^{-2}$ |
| $(rU)_{20}$ | $(1.10 \pm 0.04) \times 10^{-3}$ | $(3.5 \pm 0.1) \times 10^{-3}$ | $(4.3 \pm 0.7) \times 10^{-2}$ |
| $(rU)_{25}$ | $(2.00 \pm 0.08) \times 10^{-3}$ | $(5.2 \pm 0.1) \times 10^{-3}$ | $(1.1 \pm 0.1) \times 10^{-1}$ |
| $(rU)_{30}$ | $(2.7 \pm 0.1) \times 10^{-3}$ | $(8.1 \pm 0.8) \times 10^{-3}$ | $(2.1 \pm 0.2) \times 10^{-1}$ |
| $(rU)_{35}$ | $(4.1 \pm 0.2) \times 10^{-3}$ | $(1.1 \pm 0.1) \times 10^{-2}$ | $(4.0 \pm 1.0) \times 10^{-1}$ |
| $(rU)_{40}$ | $(4.7 \pm 0.1) \times 10^{-3}$ | $(1.30 \pm 0.06) \times 10^{-2}$ | $(6.9 \pm 0.7) \times 10^{-1}$ |
| $(rU)_{180}$ | $1.20 \pm 0.02$ | $3.2 \pm 0.6$ | $(1.0 \pm 0.2) \times 10^{2}$ |

**Supplementary Table 9. Simulation-derived dissociation constants ($K_D$) in µM.**

| **Construct** | **NTD;** $K_D$ (µM) | **RBD;** $K_D$ (µM) | **NTD-RBD;** $K_D$ (µM) |
|---|---|---|---|
| $(rU)_{10}$ | $(1.10 \pm 0.08) \times 10^4$ | 2200 ± 200 | 540 ± 10 |
| $(rU)_{12}$ | $(4.0 \pm 0.3) \times 10^3$ | 1200 ± 100 | 240 ± 9 |
| $(rU)_{15}$ | $(1.8 \pm 0.1) \times 10^3$ | 570 ± 30 | 98 ± 3 |
| $(rU)_{17}$ | $(1.20 \pm 0.08) \times 10^3$ | 400 ± 30 | 56 ± 4 |
| $(rU)_{20}$ | $(9.0 \pm 0.4) \times 10^2$ | 290 ± 10 | 24 ± 3 |
| $(rU)_{25}$ | $(5.0 \pm 0.2) \times 10^2$ | 192 ± 5 | 9.0 ± 0.6 |
| $(rU)_{30}$ | $(4.0 \pm 0.1) \times 10^2$ | 120 ± 10 | 4.7 ± 0.5 |
| $(rU)_{35}$ | $(2.4 \pm 0.1) \times 10^2$ | 92 ± 9 | 2.7 ± 0.8 |
| $(rU)_{40}$ | $(2.10 \pm 0.05) \times 10^2$ | 80 ± 4 | 1.5 ± 0.1 |
| $(rU)_{180}$ | 0.80 ± 0.02 | 0.30 ± 0.07 | $(1.0 \pm 0.1) \times 10^{-2}$ |

**Supplementary Table 10. Simulation-derived ratio of association constants $K_A$* defined as ($K_A$ of Construct + $(rU)_n$)/($K_A$ of NTD-RBD + $(rU)_{25}$).**

| **Construct** | **NTD;** $K_A$* | **RBD;** $K_A$* | **NTD-RBD;** $K_A$* |
|---|---|---|---|
| $(rU)_{10}$ | $(8.5 \pm 0.9) \times 10^{-4}$ | $(4.2 \pm 0.5) \times 10^{-3}$ | $(2.0 \pm 0.1) \times 10^{-2}$ |
| $(rU)_{12}$ | $(2.3 \pm 0.2) \times 10^{-3}$ | $(7.8 \pm 0.9) \times 10^{-3}$ | $(4.0 \pm 0.3) \times 10^{-2}$ |
| $(rU)_{15}$ | $(5.0 \pm 0.5) \times 10^{-3}$ | $(1.6 \pm 0.1) \times 10^{-2}$ | $(9.0 \pm 0.7) \times 10^{-2}$ |
| $(rU)_{17}$ | $(7.2 \pm 0.7) \times 10^{-3}$ | $(2.2 \pm 0.2) \times 10^{-2}$ | $(1.6 \pm 0.2) \times 10^{-1}$ |
| $(rU)_{20}$ | $(1.0 \pm 0.1) \times 10^{-2}$ | $(3.1 \pm 0.2) \times 10^{-2}$ | $(3.9 \pm 0.7) \times 10^{-1}$ |
| $(rU)_{25}$ | $(1.8 \pm 0.1) \times 10^{-2}$ | $(4.7 \pm 0.3) \times 10^{-2}$ | $1.0 \pm 0.1$ |
| $(rU)_{30}$ | $(2.4 \pm 0.2) \times 10^{-2}$ | $(7.3 \pm 0.9) \times 10^{-2}$ | $1.9 \pm 0.2$ |
| $(rU)_{35}$ | $(3.7 \pm 0.3) \times 10^{-2}$ | $(9.9 \pm 1.3) \times 10^{-2}$ | $3.6 \pm 0.9$ |
| $(rU)_{40}$ | $(4.2 \pm 0.3) \times 10^{-2}$ | $(1.10 \pm 0.09) \times 10^{-1}$ | $6.2 \pm 0.7$ |
| $(rU)_{180}$ | $10.7 \pm 0.7$ | $28 \pm 6$ | $900 \pm 200$ |

**Supplementary Table 11. Ion released upon binding of $(rU)_{20}$ and $(rU)_{40}$ (compare with Fig. 6 and Supplementary Fig. 5)**

| | **α (KCl)** | **α (KCl+Tris HCl)** |
|---|---|---|
| **$(rU)_{20}$** | $-3.49 \pm 0.05$ | $-5.0 \pm 0.1$ |
| **$(rU)_{40}$** | $-3.7 \pm 0.5$ | $-5.0 \pm 0.7$ |

**Supplementary Table 12. Association and dissociation constants of $NTD_L$-RBD as a function of salt concentration for $(rU)_{20}$ and $(rU)_{40}$**

| | $K_A$ ($\mu M^{-1}$) | | | | |
|---|---|---|---|---|---|
| | **50 mM KCl** | **110 mM KCl** | **150 mM KCl** | **175 mM KCl** | **200 mM KCl** |
| **$(rU)_{20}$** | 8 ± 2 | (5.4 ± 0.6) x $10^{-1}$ | (1.7 ± 0.2) x $10^{-1}$ | **-** | **-** |
| **$(rU)_{40}$** | 14 ± 2 | **-** | (3.8 ± 0.4) x $10^{-1}$ | (2.5 ± 0.3) x $10^{-1}$ | (5 ± 0.9) x $10^{-2}$ |
| | **$K_D$ ($\mu M$)** | | | | |
| | **50 mM KCl** | **110 mM KCl** | **150 mM KCl** | **175 mM KCl** | **200 mM KCl** |
| **$(rU)_{20}$** | 0.12 ± 0.03 | 1.9 ± 0.2 | 5.8 ± 0.5 | **-** | **-** |
| **$(rU)_{40}$** | 0.07 ± 0.01 | **-** | 2.6 ± 0.2 | 4.0 ± 0.4 | 19 ± 4 |

**Supplementary Table 13. $NTD_L$-RBD association constants for V21 binding**

| | $K_A$ ($\mu M^{-1}$) molecules |
|---|---|
| $K_{A1}$ | 6.2 ± 0.3 |
| $K_{A2}$ | 0.15 ± 0.10 |

**Supplementary Table 14. $NTD_L$-RBD association constants for hairpin RNA sequences**

| | $K_A$ ($\mu M^{-1}$) molecules |
|---|---|
| hpRNA | $NTD_L$-RBD |
| NSP15 | (7.8 ± 0.7) x $10^{-1}$ |
| Tetraloop | (6.7 ± 0.8) x $10^{-1}$ |
| Tetrabulge | (3.4 ± 0.7) x $10^{-1}$ |
| Decaloop | 3.4 ± 0.5 |
| SL5B | (5.3 ± 0.4) x $10^{-1}$ |

**Supplementary Table 15. Simulation-derived ratio of association constants $K_A$* defined as ($K_A$ of Construct + $(rU)_n$ )/($K_A$ of NTD-RBD + $(rU)_{25}$ )**

| **Construct** | **OmNTD-RBD (P13L,$\Delta^{31\text{-}33}$); $K_A$*** | **NTD-RBD (P13L); $K_A$*** | **NTD-RBD ($\Delta^{31\text{-}31}$); $K_A$*** |
|---|---|---|---|
| $(rU)_{25}$ | (7 ± 2) x $10^{-1}$ | 1.0 ± 0.2 | (7 ± 1) x $10^{-1}$ |
